# The PTPN1 and PTPN2 phosphatases are Cooperative Regulators of Cancer Cell Immune Evasion

**DOI:** 10.1101/2025.07.02.662801

**Authors:** Alexandre Poirier, Erika Walback, Rui Su, Isabelle Aubry, Chenyue Wu, Elisabeth St-Laurent, Sonali Uttam, Ana Maria Hincapie, Bianca Colalillo, Serge Hardy, Samuel Doré, Michel L. Tremblay

## Abstract

Immune evasion by cancer cells remains a major barrier to the success of immune checkpoint blockade (ICB). Here, we identify the phosphatases PTPN1 and PTPN2 as cooperative regulators of tumor immune resistance. Dual genetic ablation of PTPN1/2 in cancer cells enhances Type I and II interferon signaling, MHC-I and CXCL9 expression, and sensitizes tumor cells to cytotoxic T lymphocyte–mediated killing. The small-molecule inhibitor KQ791 phenocopies these effects and synergizes with anti-PD1 therapy to suppress tumor growth in murine models, including immunotherapy-refractory cancers. Mechanistically, PTPN1/2 loss augments STAT1/3/5 signaling and primes cancer cells for immunogenic cell death via IFNγ/TNFα-induced pathways. Moreover, PTPN1/2 inhibition enhances antigen release and cross-presentation, promoting robust antigen-specific CD8⁺ T cell responses. These findings highlight PTPN1&2 as essential mediators of cancer immune evasion and support their inhibition as a strategy to broaden the effectiveness of immune checkpoint blockade in solid tumors.

## Introduction

Immune checkpoint blockade (ICB) has revolutionized the way we treat patients suffering from cancer. However, many patients remain refractory to immunotherapy, highlighting the importance of identifying new genetic targets that can achieve therapeutic outcomes [1–3]. One such target is the protein tyrosine phosphatase non-receptor 2 (PTPN2, aka T-cell phosphatase or TC-PTP). Indeed, the deletion of PTPN2 enhances T cell immunity against cancer by increasing T cell receptor (TCR) signaling [4], cytokine responses [5–7] and proliferation [8]. Unlike most immunotherapy targets, PTPN2 is also targetable with small-molecule inhibitors [9, 10]. Our group and others have demonstrated that targeting PTPN2 with small-molecule inhibitors enhances anti-cancer immune responses [10–13]. Mechanistically, PTPN2 inhibition boosts the activity and fitness of cytotoxic T lymphocytes (CTLs) by increasing the activity of the STAT5 and IL-2 signaling pathways [10, 13]. Competitive inhibitors of PTPN2 also antagonize PTPN1 (also known as PTP1B) at equimolar concentrations due to high structural resemblance in their respective catalytic sites [13, 14]. For enhancing the activity of T cells, we have shown that dual deletion of PTPN1 and PTPN2 is favorable over deletion of PTPN2 alone, thus mitigating initial concerns over the lack of specificity of PTPN2 inhibitors [12].

The two phosphatases exhibit overlapping roles in inhibiting the JAK/STAT signaling pathway in response to various cytokines and interferons. PTPN1 preferentially suppresses JAK2 and TYK2 kinases, whilst PTPN2 suppresses more potently JAK1 and JAK3 kinases and can directly inhibit STAT1 and STAT3 transcription factors [15–18]. The two enzymes also have non-overlapping roles due to differences in substrates and subcellular localization [19–22]. PTPN1, which is tethered to the endoplasmic reticulum, can dephosphorylate members of the receptor tyrosine kinase family (RTKs), such as the insulin receptor (IR), EGFR, and MET [20, 23–26]. PTPN1 also opposes oncogenic RAS signaling by upregulating the GTPase activating protein RASA1 (p120RasGap) [27]. In cancer cells, PTPN1 inhibition has been shown to limit the proliferation of breast cancer cells via its role in HER2/ErbB2 and STAT5 signaling [28–32]. PTPN1 also facilitates the survival of breast cancer cells in hypoxic conditions by suppressing inflammatory cell death programs [33, 34]. Conversely, PTPN2 ablation has been found to improve immune responses against breast cancer via T cell and tumor cell intrinsic mechanisms [7]. The capacity of melanoma-initiating cells to evade immune recognition has also been linked to PTPN2 via its suppression of STAT1 [35]. Still, it remains unclear whether inhibition or deletions of these phosphatases could affect cancer cell immune evasion. Given that a phase I cancer immunotherapy clinical trial using a PTPN2/1 inhibitor is ongoing (<u>NCT04777994</u>), understanding the potential impact of inhibition in tumor cells is essential.

Herein, we found that the two phosphatases act synergistically to suppress Type I interferon programs in cancer cells, thereby inhibiting their recognition by CTLs. Furthermore, we found that pharmacological inhibition of PTPN1/2 sensitized multiple solid tumor models to the effects of anti-PD1 immunotherapy, in a cancer cell autonomous fashion. Ablation of Ptpn1/2 directly in cancer cells enhanced the establishment of cellular immunity against tumors by promoting immunogenic cell death through a JAK/STAT signaling axis. Therefore, the enzymatic activity of PTPN1/2 promotes cancer cell immune evasion by decreasing the visibility, targetability, and cell death of cancer cells. PTPN1/2 inhibition thus complements PD1 blockade, which could expand its use to more patients and cancer types.

## Results

### Ptpn1&2 synergistically restrict the recognition and killing of cancer cells by cytotoxic T lymphocytes

To understand the possible contribution of Ptpn1 and Ptpn2 in mediating cancer cell immune evasion, we first knocked down these two phosphatases in B16-F10 melanoma cells that constitutively express the chicken Ovalbumin protein (B16-OVA) (Extended Data Fig. 1a). Chicken ovalbumin (OVA) served as the model antigen in T cell cytotoxicity assays. B16-OVA cells transfected with the different siRNAs were co-cultured with OT-1 T cells, which express a T cell receptor (TCR) restricted to the OVA_257-264_ peptide [36]. Knocking down both Ptpn1 and Ptpn2 enhanced the capacity of OT-1 to kill B16-OVA cells, whilst single knockout conditions had no significant effect (Fig. 1a and b). Additionally, since Ptpn1/2 have a putative role in suppressing Type I and II interferon signaling [15, 18, 37], B16 OVA cells were also stimulated with IFNβ and IFNγ for 24 hours, which causes the transcription of MHCI genes. We then quantified the expression of MHCI (H2KB in mouse) on the surface of B16-OVA cells with the various knockdown conditions. We found that dual knockdown resulted in the most prominent increase in MHCI (H2KB) expression, either in unstimulated or in IFNβ/IFNγ-stimulated B16-OVA cells (Fig. 1c, Extended Data Fig. 1b). Knockdown of Ptpn1 alone also enhanced MHCI expression in unstimulated and IFN-stimulated cells, whilst Ptpn2 knockdown only led to a slight enhancement compared to siCTL-treated cells (Fig. 1c, Extended Data Fig. 1b). We next assessed whether the knockdown of Ptpn1/2 enhanced the secretion of cytokines by cancer cells in response to IFNγ using a mouse chemokine array (Fig. 1d, Extended Data Fig. 1). To enhance the production of secreted factors, cells were stimulated with IFNγ for 24 hours. We used IFNγ rather than IFNβ since the latter is better established as a factor leading to pro-inflammatory cytokine/chemokine release in non-immune cells [38, 39]. We observed a synergistic enhancement of Cxcl1, Cxcl9, Cxcl10 and Ccl2 secretion B16-OVA cells with dual Ptpn1/2 knockdown (Fig. 1d, Extended Data Fig. 1). Out of all the detected factors, Cxcl9 showed the most prominent enhancement in siPtpn1&2 treated cells, with a more than 5-fold increase following IFNγ stimulation (Fig. 1d, Extended Data Fig. 1). In comparison, siPtpn2 treated cells only had an increase of 66% in Cxcl9 levels versus siCTL cells when treated with IFNγ (Fig. 1d, Extended Data Fig. 1). Cxcl9 is a chemokine that binds to the Cxcr3 receptor on effector T cells and leads to their recruitment to sites of infection and tumors [40]. We validated the synergistic enhancement of Cxcl9 in siPtpn1/2 transfected cells using flow cytometry (Extended Data Fig. 1d). The frequency of Cxcl9-positive cells raised from 7.5% in siCTL treated cells to 77% in siPtpn1&2 treated cells, 24 hours after the addition of IFNγ (Extended Data Fig. 1d). Thus, the ablation of both these phosphatases in target cells leads to a synergistic enhancement in cytolysis, MHCI expression, and chemokine secretion, most notably Cxcl9.

**Figure 1:**
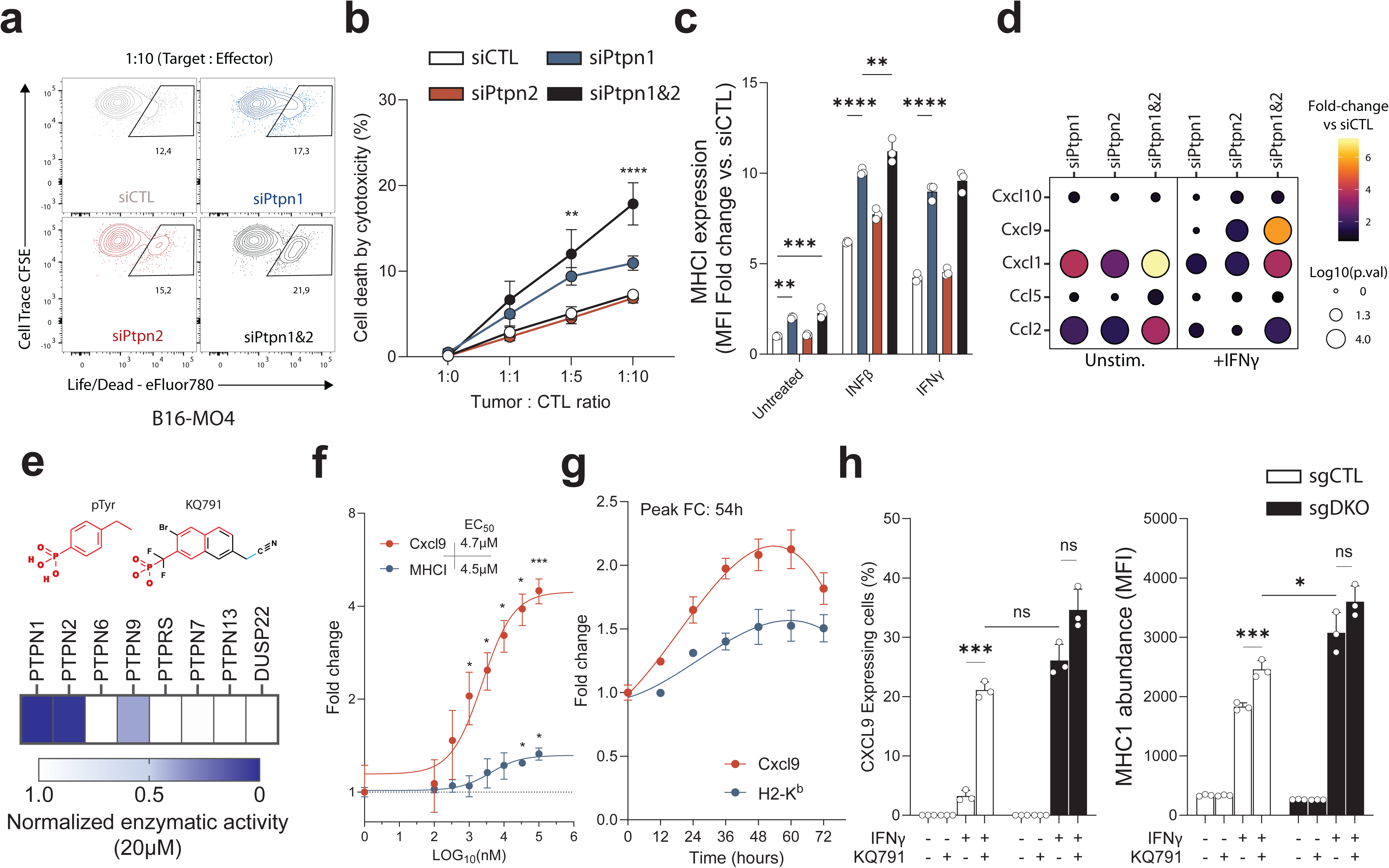
Dual, but not single, targeting of Ptpn1 and Ptpn2 sensitizes cancer cells to T cell cytolysis. a) Flow cytometric analysis of the viability of B16-F10-OVA cells with the indicated knockdown conditions after co-culture with OT-1 T cells. B16-OVA cells were labeled with cell trace CFSE prior to co-culture to discern between cancer and T cell viability. b) Frequency of cell death in B16-OVA cells co-cultured with increasing amounts of OT-1 cytotoxic T lymphocytes (CTL). c) Fold change in mean fluorescence intensity (MFI) of mouse MHCI (H2KB) with Ptpn1, n2 or dual knockdown and IFNβ or IFNγ treatment measured by flow cytometry. Fold change is relative to siCTL untreated cells. d) Relative chemokine abundance in B16-OVA cells with the indicated knockdowns and stimulated with or without IFNγ for 24h. Chemokine abundance is normalized to either unstimulated siCTL (left) or IFNγ-treated siCTL cells (right). e) Molecular structures of phospho-tyrosine (pTyr) and of the PTPN1/2 dual inhibitor KQ791 (top). Normalized enzymatic activity of human PTPN1, PTPN2, and other phylogenically related tyrosine phosphatases upon treatment with 20μM of KQ791 (bottom). f) Fold change in Cxcl9 and MHCI (H2KB) MFI in B16-OVA upon treatment with IFNγ and escalating concentrations of KQ791 measured by flow cytometry. Fold change is relative to cells treated with IFNγ alone. g) Fold change in Cxcl9 and MHCI (H2KB) MFI in B16-OVA cells after treatment with IFNγ for 24 hours and KQ791 for the indicated times. Fold change is relative to cells treated with IFNγ alone. h) Percentage of Cxcl9 expressing cells (Left) and the MFI of MHCI (H2KB) (Right) in control (sgCTL) or double knockout (sgDKO) treated with or without IFNγ and/or KQ791 (20μM). Data pooled from two independent experiments (a and b). c, e, f, g, and h) Data representative of three independent experiments done in triplicate (c, e, f, g, and h). Error bars represent mean±s.e.m. Statistical analysis: Two-way ANOVA with Tukey’s multiple comparisons. Key: ns nonsignificant, *p<0.05, **p<0.01, ***p<0.001 and ****p<0.0001.

### KQ791, a small-molecule PTPN1/2 dual inhibitor, phenocopies dual deficiency in B16-OVA cells

To further investigate the role of Ptpn1 and Ptpn2 in cancer cell immune evasion, we first sought to validate whether pharmacological inhibition could phenocopy the effects observed in Ptpn1/2 deficient B16-OVA cells. We tested L598 and its optimized derivative, KQ791, two small-molecule dual inhibitors with nanomolar efficiency for human PTPN1 and PTPN2 (Extended Data Fig. 1e and f). These inhibitors were initially designed to be orally bioavailable insulin sensitizers for the treatment of type 2 diabetes [41–45]. Although deemed safe and tolerable in humans, KQ791 failed at the clinical stage due to a lack of efficacy against diabetes (1a: <u>NCT02445911</u> and phase IB/II <u>NCT02370043</u>). Thus, we aimed to repurpose this molecule as a tool compound for cancer immunotherapy. In enzymatic and cellular assays, KQ791 was found to have enhanced activity versus L598 regarding target inhibition and Cxcl9 release (Extended Data Fig. 1f and g). We thus pursued our investigation with KQ791 and confirmed its specificity over closely related phosphatases such as PTPN6 (SHP-1), PTPRS (PTP-σ), PTPN7 (HePTP), PTPN13 (PTPL1), PTPN9 (MEG2), and one unrelated dual specificity phosphatase DUSP22 (Fig. 1e, Extended Data Fig. 1h). Although KQ791 also partially inhibited PTPN9 at a high concentration, we observed a 37.7-fold difference in the IC_50_ of PTPN1 over PTPN9 with respective IC_50_ of 188nM and 7199nM (Extended Data Fig. 1i). In cellular assays, KQ791 exhibited a concentration-response behavior leading to an approximate 400% and 40% enhancement in Cxcl9 and MHCI abundance after IFNγ stimulation (Fig. 1f). In these assays, KQ791 had an observed cellular effective concentration 50% (EC_50_) of approximately 4.5μM, with a plateaued effect in concentrations above 30μM (Fig. 1f). The peak response following treatment with KQ791 was also found to be around 54-hours, indicating a significant delay in the mounting of response to dual inhibitors (Fig. 1g). Finaly, we sought to validate whether KQ791 could phenocopy the effects of dual deficiency in terms of Cxcl9 and MHCI expression. To do so, we generated CRISPR/Cas-mediated Control, Ptpn1, Ptpn2, and Ptpn1+Ptpn2 (DKO) knockout B16-OVA cell lines (hereafter sgCTL, sgPtpn1, sgPtpn2, and sgDKO)(Extended Data Fig. 1j and k). Stable knockout cells represented a more robust tool for our investigations compared to our initial approach with siRNAs. Treatment with KQ791 and IFNγ in sgCTL cells had a similar effect to stimulation with IFNγ alone in sgDKO cells regarding Cxcl9 and MHCI abundance (Fig. 1 h). Furthermore, the addition of KQ791 to sgDKO cells did not result in a significant increase in the expression of these proteins, suggesting that the effect of KQ791 is mediated only by PTPN1/2 (Fig. 1h). Overall, these observations indicate that the use of a small-molecule inhibitor can recapitulate the previous effects observed with PTPN1/2 genetic deletion in cancer cells and therefore that the catalytic activity of these phosphatases is likely responsible for these phenotypes.

### Multi-omics analysis of Ptpn1/2 deficient cells reveals the formation of a spontaneous Type I interferon response

In parallel with the initial experiments, we performed proteomics analyses on siCTL or siPtpn1&2-treated B16-OVA cells to explore global changes in cellular responses. We found that knockdown of these two phosphatases increased the protein abundance of several genes related to the response to Type I interferons (IFNα/β) and to glycolysis using gene enrichment analysis (Fig. 2a). We observed a more than 6-fold increase in the Type I interferon response genes Tap1, Ifit2, Ifit3, Ifit3b and Ifi204 in unstimulated double knockdown cells compared to control (Fig. 2a). Extracellular Acidification Rate (ECAR) quantifications with Seahorse on B16-OVA cells with the various genotypes confirmed a slight enhancement of glycolysis in the absence of Ptpn1/2 (Extended Data Fig. 2a and b). To assess the transcriptional effect of deleting each phosphatase alone or conjunctly, we sent isolated RNAs from sgCTL, sgPtpn1, sgPtpn2, sgDKO, and sgCTL treated with KQ791 for bulk RNA sequencing. Additional samples from the same conditions were stimulated with IFNβ for 6 hours before RNA isolation and sequencing, given that we had observed this signature in the Ptpn1/2 knockdown cells. Consistent with our previous results, analysis of the gene ontologies from the differentially expressed genes (DEGs) also revealed the presence of a Type I interferon signature in Ptpn1/2 deficient unstimulated B16-OVA cells (Fig. 2b). sgPtpn1, sgPtpn2, and KQ791-treated cells were also found to have an enhancement of this signature, albeit to a lesser extent than sgDKO cells (Fig. 2b). We also detected the presence of an IFNγ response signature in sgPtpn2 and a signature of the extrinsic apoptotic pathway in sgPtpn1 cells (Fig. 2b). Both these signatures were carried over in sgDKO cells (Fig. 2b). To then deconvolute how the knockout of each phosphatase led to the differential expression of specific genes, we performed an intersecting set analysis and visualized it using upset plots (Fig. 2c). All conditions had increased abundance of *Ccl2* and *Isg15*, a canonical IFN-response gene (Fig. 2c). On the one hand, *Cxcl9* was only found to be upregulated in sgPtpn2, sgDKO, and KQ791-treated cells, (Fig. 2c). indicating that Ptpn2 blockade was responsible for the increase in *Cxcl9*. Conversely, Ptpn1 ablation alone did not enhance the transcription of *Cxcl9 but* was found to be linked to the increase in *Trail* and *Tnfrsf9* (Fig. 2c). We next explored the DEGs in response to treatment with IFNβ and focused on three gene sets that were highly enriched in sgDKO cells: response to IFN, TNFR1 signaling, and pattern recognition receptors (PRRs) and their related effectors (Fig. 2d). In the majority of cases, DEGs from these three sets were increased/decreased the most in sgDKO (Fig. 2d). sgPtpn2 cells also had a substantial increase in these DEGs, whilst sgPtpn1 cells did not (Fig. 2d). These observations thus suggest a functional dominance of Ptpn2 over Ptpn1 in restricting the response to IFNβ. In an intersection analysis on the samples stimulated with IFNβ, we observed that the vast majority of DEGs from the sgDKO cells were shared with sgPtpn2 (*Irf1*, *Cd274*, *Ripk1*) (Extended Data Fig. 2c). We also observed the upregulation of the checkpoint ligand *Pdl1* (*Cd274*) in both unstimulated and IFNβ-stimulated sgPtpn2 and sgDKO cells (Extended Data Fig. 2d). We next intersected the DEGs in IFNβ-KQ791-treated cells with the other genotypes to understand how well the inhibitor was recapitulating DEG diversity (Fig. 2e). In line with previous observations, we found that KQ791-treated cells shared 76%, 75%, and only 25% of their DEGs with sgPtpn2, sgDKO, and sgPtpn1 cells, respectively (Fig. 2e). Although the degree to which each of theses DEGs was increased/decreased was not as substantial as the genetic knockouts, this result suggests that the inhibitor most closely recapitulates the phenotype of sgDKO/sgPtpn2 cells (Extended Data Fig. 2e). Given the large number of DEGs observed in IFNβ-treated sgDKO cells, we sought to compare the DEGs in stimulated controls versus stimulated sgDKOs (Fig. 2f, Extended Data Fig. 2e and f). We found that among the 1733 DEGs found in sgDKO + IFNβ, less than half (817) were shared with sgCTL + IFNβ cells (Fig. 2f). These genes, termed canonical IFNβ-response genes, included *Cxcl9*, *Pdl1* (*Cd274*), and *Isg15* (Fig. 2f). On the other hand, we found 916 DEGs unique to sgDKO-stimulated cells, such as *Ripk1*, *Sting1*, *Nos2*, and *Socs3* (Fig. 2f). To understand what the signaling cascades and transcription factors (TFs) linked to this difference were, we performed TF activity analyses on sgDKO-stimulated cells using decoupleR and collecTRI [46]. This analysis revealed a significant increase in Stat1, Stat3, Irf1, Nfkb, and Stat5 TF activity, among others (Fig. 2g). These data suggest that Ptpn1/2 ablation enhances the response to IFNs and generates a singular response that coincides with enhanced cytolysis by T cells. To validate this observation functionally, we repeated in vitro killing assays using sgCTL and sgDKO cells, either pre-emptively stimulated with IFNβ or not (Fig. 2h). The addition of IFNβ in sgCTL cells resulted in a degree of cytolysis that was similar to that of unstimulated sgDKO cells, both around 15% in the highest Target:CTL ratio (Fig. 2f). Addition of IFNβ to sgDKO cells further increased cytolysis to around 40%, surpassing the controls by more than 1-fold (Fig. 2f). Dual ablation of Ptpn1/2 thus results in a synergistic enhancement of Type I interferon responses and MHC-class I expression (Extended Data Fig. 2f), which sensitizes cancer cells to CTL-mediated killing.

**Figure 2:**
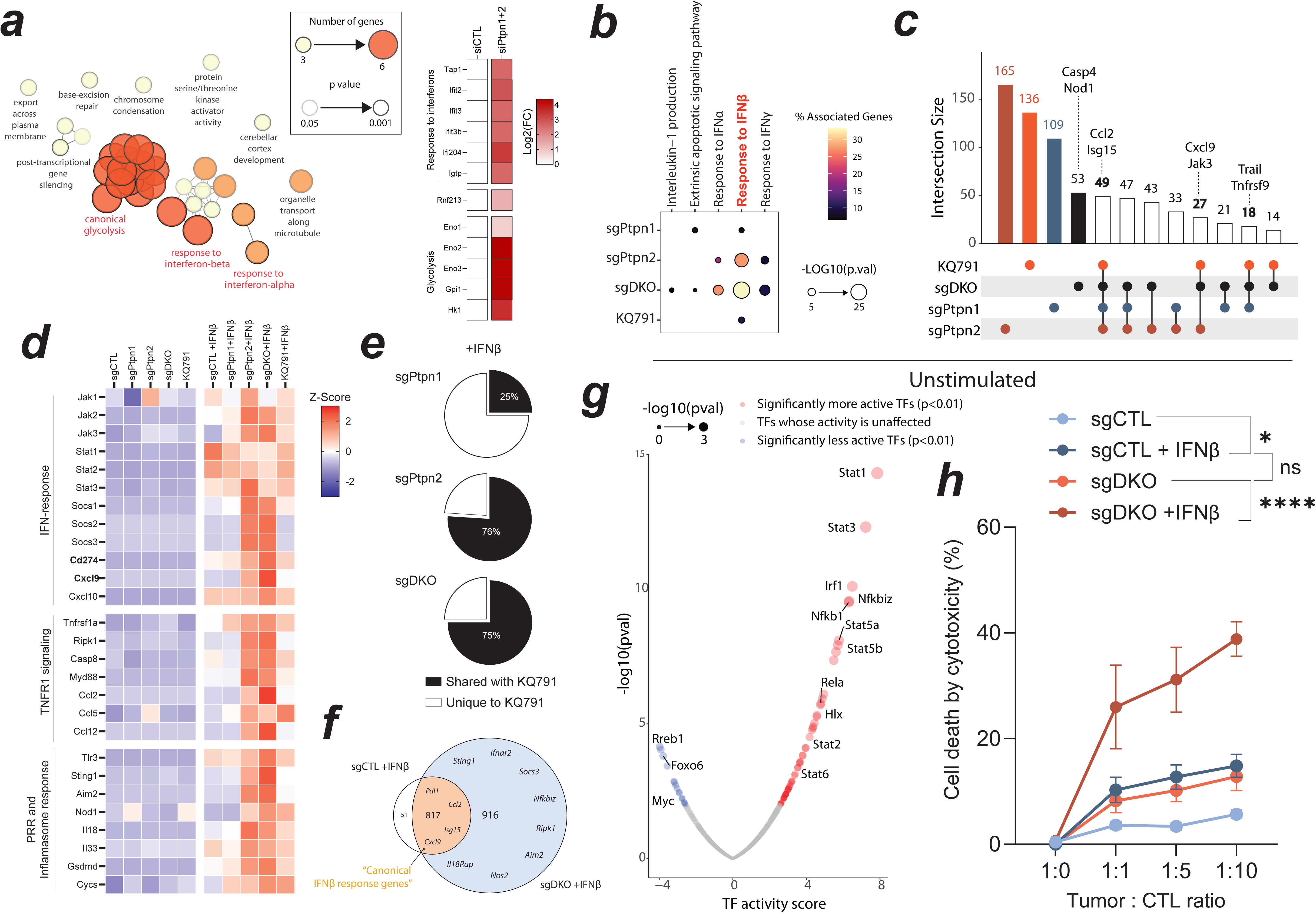
Dual ablation of Ptpn1&2 induces a Type I interferon response in cancer cells. a) Top gene ontologies from a comparative proteomic analysis in unstimulated B16-OVA cells with Ptpn1&2 dual knockdown versus control siRNA (siCTL) (left). Heatmap of the Log2(Fold-change) of the proteins of interest with increased abundance in siPtpn1&2 versus siCTL cells (right). b-g) Bulk RNAseq analysis of B16-OVA cells with stable knockout of either Ptpn1, Ptpn2, Ptpn1&2 (sgDKO), control or control treated with KQ791 (20μM for 24h). b) Bubble plot of the significantly over-represented gene ontologies in the different genotypes (Untreated cells). c) Upset plot of the differentially expressed genes (DEGs) that are common between the indicated sets of conditions (Untreated cells). Genes of interest related to the ontologies in b) are labeled above the respective conditions that share these DEGs. d) Z-score heatmap for DEGs belonging to the IFN response, TNFR1, or PRR and inflammasome gene clusters. Untimulated cells are shown on the left side of the heatmap, while cells treated with IFNβ are shown on the right. Selected DEGs are significant (FDR < 5%) in at least one of the conditions. e) Pie chart of the frequency of DEGs (LOG2(FC) >1 or <-1, FDR<1%) shared between KQ791 treated cells and sgPtpn1, sgPtpn2 or sgDKO cells. All comparisons are based on the list of DEGs significantly different from sgCTL cells treated with IFNβ. f) Venn diagram of the number of DEGs in the sgDKO + IFNβ that are common or unique to sgCTL + IFNβ. The list of DEGs in each condition was compared to sgCTL unstimulated cells. g) Volcano plot of the transcription factor (TFs) activity score in sgDKO + IFNβ versus sgCTL + IFNβ B16-OVA cells. TFs with significantly enhanced activity scores are labeled in red, while those with decreased activity are shown in blue. h) Frequency of cell death in B16-OVA cells co-cultured with increasing amounts of OT-1 CTLs. sgCTL or sgDKO cells were either pre-treated with IFNβ or left unstimulated before co-culture with OT-1 CTLs. b-g) Bulk RNAseq was performed on three independent replicates for each condition. N=30 samples. H) Data pooled from two independent experiments done in triplicate. Error bars represent mean±s.e.m. Statistical analysis: h) Two-way ANOVA with Šídák’s multiple comparisons test. Key: ns nonsignificant, *p<0.05, **p<0.01, ***p<0.001 and ****p<0.0001.

### Ptpn1/2 ablation or inhibition increases Stat1, 3, and 5 activities in response to IFNγ

Next, we set our sights on characterizing the biochemical responses of Ptpn1/2 knockout cells. Although we initially focused on Type I interferon responses, Type II interferons (IFNγ) are known to be particularly relevant in cancer immunotherapy. When comparing IFNβ to IFNγ stimulations side-by-side in B16-OVA cells, we found that only the latter could induce the production of Cxcl9 in sgCTL cells (Extended Data Fig. 2a). We then proceeded to stimulate the various B16-OVA cell lines with IFNγ for 30 minutes and analysed the amount of phosphorylation of Stat1, Stat3, and Stat5 using western blotting (Fig. 3a). We focused on these three putative substrates since they were found to be the most active in the absence of Ptpn1/2 (Fig. 2g). Stat1 phosphorylation (Tyr701) was only found to be substantially augmented in sgPtpn2, sgDKO, and KQ791-treated cells (Fig. 3a). We observed an increase in Stat3 and Stat5 phosphorylation (Tyr 705 and 694, respectively) in both single and dual ablation of Ptpn1/2, with an apparent synergistic effect on Stat5 phosphorylation in sgDKO and KQ791-treated cells (Fig. 3 a). We also observed basal hyperphosphorylation of Stat1 and Stat3 in sgDKO cells (Fig. 3a; long exposure). These results suggest that among these two phosphatases, Ptpn2 is the most critical negative regulator of Stat1 activation, while Ptpn1 and Ptpn2 conjunctly suppress Stat3 and Stat5 phosphorylation.

**Figure 3:**
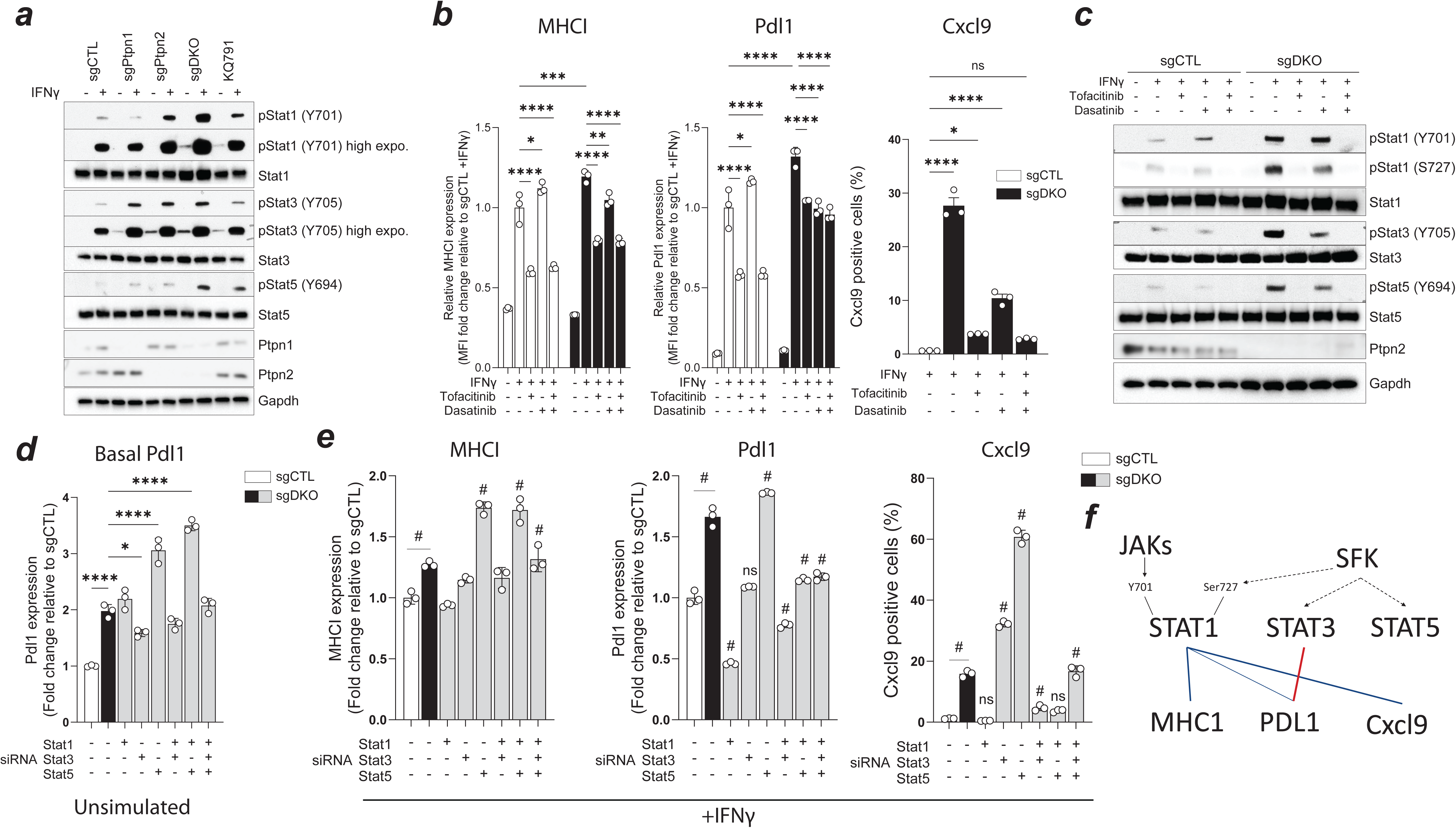
Heightened response to interferon relies on both JAK/STAT signaling and Src-family kinase activity. a) Western blot of phosphorylated and total STAT1,3 and 5 in B16-OVA cells treated with IFNγ for 30 minutes or left untreated. Gapdh is used as the loading control. b) Flow cytometric analysis of MHCI (H2KB), Pdl1, and Cxcl9 expression in sgCTL and sgDKO cells treated with IFNγ, Tofacitinib (1μM) and/or Dasatinib (1μM). Data was normalized to sgCTL-treated cells (+IFNγ) for MHCI and Pdl1. Tofacitinib is a JAK1-3 inhibitor. Dasatinib is an ABL1/2 and Src-family kinase inhibitor. c) Western blot of phosphorylated and total STAT1,3 and 5 in B16-OVA cells treated with IFNγ and co-treated with Tofacitinib (1μM) and/or Dasatinib (1μM). Gapdh is used as the loading control. d) Flow cytometric analysis of Pdl1 expression in unstimulated sgCTL or sgDKO cells transfected with siRNAs for Stat1,3 and 5. Data was normalized to sgCTL cells. e) Flow cytometric analysis of MHCI (H2KB), Pdl1, and Cxcl9 expression in sgCTL and sgDKO cells treated with IFNγ and transfected with siRNAs for Stat1,3 and 5. Data was normalized to sgCTL-treated cells (+IFNγ) for MHCI and Pdl1. f) Relationship graph for the observations reported in Fig. 3. Complete lines represent a direct relationship between the reported markers. Hashed lines represent an indirect relationship between Src-family kinases (SFKs) and STAT1,3 and 5. Data representative of at least three independent experiments. Error bars represent mean±s.e.m. Statistical analysis: Two-way ANOVA with Tukey’s multiple comparisons test (b[MHCI, Pdl1]). One-way ANOVA with Dunnett’s multiple comparison test (b[Cxcl9], d, and e). Key: ns nonsignificant, *p<0.05, **p<0.01, ***p<0.001 and ****p<0.0001. In e), “#” denotes significance of at least p<0.05 versus the sgCTL condition (white bar).

### Src-family kinase activity provides accessory signaling to Stats in the absence of Ptpn1 and Ptpn2

The JAK/STAT pathways are regulated by numerous tyrosine kinases that operate at distinct levels in the pathway. This includes the canonical activators, JAK1-3 (JAKs) and TYK2, but also members of the Src family of kinases (SFKs) such as SRC, ABL1/2, and FYN, which are also substrates of PTPN1/2 [30, 47, 48]. Given the contribution of two families of kinases to the activation of STATs, we sought to understand the relevance of each family during IFNγ signaling. We performed cellular assays on IFNγ-stimulated B16-OVA cells with or without concomitant inhibition of JAKs (Tofacitinib) and/or SFKs (Dasatinib). JAKs inhibition with Tofacitinib led to a sharp decrease in MHCI (H2KB), Pdl1, and Cxcl9 expression (Fig. 3b, Extended Data Fig. 3b). Unlike sgCTL cells, sgDKOs were found to have a decrease in Pdl1 and Cxl9 expression following SFKs inhibition with Dasatinib (Fig. 3 b, Extended Data Fig. 3 b). Most notably, both Tofacinib and Dasatinib were required to decrease Cxcl9 expression and return it to control levels in sgDKOs (Fig. 3 b, Extended Data Fig. 3 b). Western blotting analyses revealed that treatment with Dasatinib affected Stat3 and Stat5 tyrosine phosphorylation but had no effect on Stat1 (Tyr 701) (Fig. 3c). However, the phosphorylation of another essential residue for Stat1 activity, Ser 727, was found to be decreased by SFKs inhibition (Fig. 3c)[49]. These data suggest that the activity of SFKs is providing an accessory signal for Stat activation in the absence of Ptpn1/2. Given that SFK inhibition affected Stat1, 3, and 5 differently, we sought to understand the importance of each of these TFs on MHCI, Pdl1, and Cxcl9 expression, using STATs siRNAs (Fig. 3 d and e, Extended Data Fig. 3c and d). Without IFNγ stimulation, Stat3 appeared to be related to the increase of Pdl1 expression observed previously in sgDKO cells (Fig. 3d). During stimulation, however, MHCI, Pdl1, and Cxcl9 were most drastically decreased by knockdown of Stat1 (Fig. 3e). We observed a compensation of Stat1 protein levels following Stat3 and Stat5 knockdown, making the specific contributions of Stat3 and Stat5 difficult to interpret (Extended Data Fig. 3c). Nonetheless, these observations reveal the importance of SFKs, most likely through the indirect activation of Stat1 Ser 727, in augmenting antigen presentation and Cxcl9 release in Ptpn1/2 deficient tumor cells (Fig. 3f).

### Identifying the cell lines and cancer types that are the most sensitive to PTPN1/2 inhibition

Before testing the efficacy of KQ791 on solid tumors in vivo, we sought to screen different human cell lines to assess which cancer type could be the most sensitive to PTPN1/2 inhibition. We cultured MiaPaca2 (Pancreatic), MCF-7 (Breast), OVCAR-3 (Ovarian), U251 (Glioblastoma), U-87-MG (Glioblastoma), A549 (Lung), HCT116 (Colon), Hela 299 (Cervical), A431 (Squamous cell carcinoma) and LnCap (Prostate) cancer cell lines and stimulated them with IFNγ, with or without KQ791 co-incubation (Fig. 4 a-d). We found that among these cell lines, HCT116, MCF-7, MiaPaca2, and U251 had the most robust increase in MHCI (HLA A, B, C), PDL1, and CXCL9 in response to IFNγ and KQ791 co-stimulation (Fig. 4a-c, Extended Data Fig. 4a and b). We also observed a modest decrease in the proliferation rates of these cell lines in response to KQ791 treatment with IFNγ (Fig. 4d). These data suggest that cell lines derived from colon and breast cancers could be more responsive to PTPN1/2 inhibition. To understand the molecular determinants of the response, we correlated the amplitude of CXCL9 secretion to the transcriptome of all the cell lines screened in this assay using the Human Protein Atlas Cancer Cell line Dataset (Fig. 4e and f) [50]. We identified several genes of the Interleukin 1 family (IL1B, IL16, IL33, IL36A/B) along with some SFKs (FYN, BLK) to be positively correlated to the enhancement in CXCL9 secretion following PTPN1/2 inhibition (Fig. 4e). Further analysis of the TFs associated with the response revealed RFXAP, RELA, NFKB, SPI1, STAT1, and IRF1 among the top factors related to the enhanced sensitivity to KQ791. Overall, these analyses suggest that cell lines derived from solid cancers that express IL1-family cytokines and that have high NFKB, STAT1, and IRF1 activity are the most predisposed to respond to PTPN1/2 inhibition.

**Figure 4:**
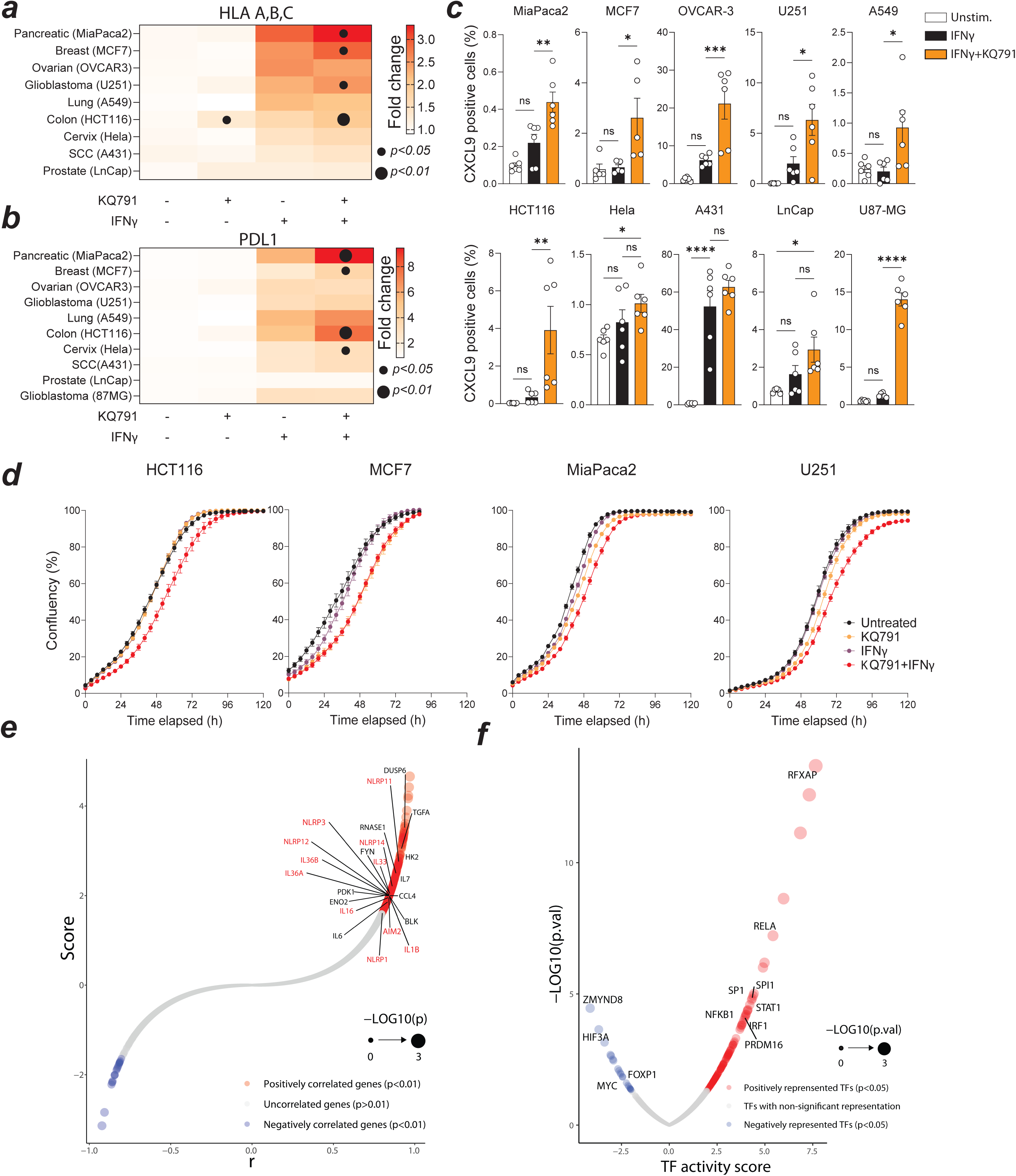
Analysis of the effect of KQ791 in various human cancer cell lines identifies HCT116 colon and MCF-7 breast cancers as top responders. Flow cytometric analysis of the MFI fold-change in HLA,A,B,C (a) and PDL1 (b) relative to unstimulated, untreated cells. c) Frequency of CXCL9-positive cells in the various human cancer cell lines after treatment with IFNγ and KQ791 measured by flow cytometry. d) Proliferation curves for the top 4 cancer cell lines from a-c). Cells were incubated with or without IFNγ and KQ791 and left to proliferate in an Incucyte apparatus for up to 120 hours. e) Bubble plot of the gene transcripts that are either positively (red) or negatively (blue) correlated to the response to CXCL9 from c). The correlation score was used to rank the top genes and was calculated by multiplying the p-value and the Pearson correlation coefficient (r). Genes labeled in red are genes related to the inflammasome response and the IL1 family of cytokines. Transcript abundance from the human cell lines was obtained from the Human Protein Atlas server. f) Analysis of Transcription factor (TFs) activity from the dataset generated in e). TFs with significantly represented activity scores are labeled in red, whilst those with decreased representation are shown in blue. Data pooled from two or three independent experiments done in triplicate (a-c). Data representative of three independent experiments (d). Error bars represent mean±s.e.m. Statistical analysis: Two-way ANOVA with Tukey’s multiple comparisons (a and b). One-way ANOVA with Dunnett’s multiple comparison test (c). Pearson correlation analysis between transcript abundance and the fold change in CXCL9 positivity between IFNγ and IFNγ + KQ791 treated cells (e). Key: ns nonsignificant, *p<0.05, **p<0.01, ***p<0.001 and ****p<0.0001.

### Ptpn1/2 dual inhibition sensitizes solid tumors to checkpoint blockade in syngeneic models that are either sensitive or insensitive to anti-PD1

We next investigated the capacity of KQ791 to induce anti-tumoral immune responses in syngeneic cancer models. We first tested the compound in MC38 (Colon cancer model) and EMT-6 (triple-negative breast cancer model) tumors, given that we had previously identified these tumor types as most sensitive to PTPN1/2 inhibition in our previous experiments (Fig. 5a-e). We sought to compare the efficacy of PTPN1/2 inhibition to the standard of care immunotherapy, anti-PD1. We included a group that received both treatments (KQ791 and anti-PD1 combo), given that we had observed an upregulation of PDL1 in cancer cells following PTPN1/2 inhibition or deletion. To reduce potential biases, these two initial trials were conducted by an independent contractual research organization (CRO), where groups were anonymized and randomized upon treatment. We found MC38 tumors to be sensitive to anti-PD1, but not to KQ791 single treatments (Fig. 5a, Extended Data Fig. 5a). Combination treatment resulted in the most significant reduction in tumor growth among all the groups (Fig. 5a, Extended Data Fig. 5a). To better understand the differences in therapeutic efficacy, we characterized the immune populations within tumors using spectral flow cytometry (Fig. 5b-d). We found a significant increase in the total frequency of Cd45+ immune cells in tumors treated with anti-PD1 and Combo groups (Fig. 5b). Tumors treated only with KQ791 had only a modest increase in the frequency of immune cells (p=0.054). Further inspection of the major subsets of immune cells within the Cd45+ population revealed an increased proportion of Cd8+ T cells in MC38 tumors treated with KQ791, anti-PD1, and Combo therapy (Fig. 5c). We found that Cd8+ T cells from the Combo group had a decreased expression of Cd62l (a naïve T cell marker) and increased expression of the effector markers Cd44 and Cxcr3 (Cxcr3 being the receptor for Cxcl9) (Fig. 5d). These data suggest that the addition of KQ791 with anti-PD1 either enhances the trafficking of effector T cells to tumors or facilitates the transition of T cells to effector cells within tumors. In the second model tested, EMT-6 triple negative breast cancer (TNBC) tumors, we found that none of the single arm groups had a meaningful effect on tumor growth (Fig. 5e, Extended Data Fig. 5 b). However, the Combo therapy was found to have a substantial impact, both on tumor burden and in survival of EMT-6 bearing mice (Mean growth inhibition of 72% and absence of visible tumors in 7 of 12 animals at day 40) (Fig. 5e, Extended Data Fig. 5c). Since KQ791 sensitized the effect of anti-PD1 in a non-responsive model, we performed a third trial in the LLC1 lung cancer model, which is insensitive to immunotherapy [51].

**Figure 5:**
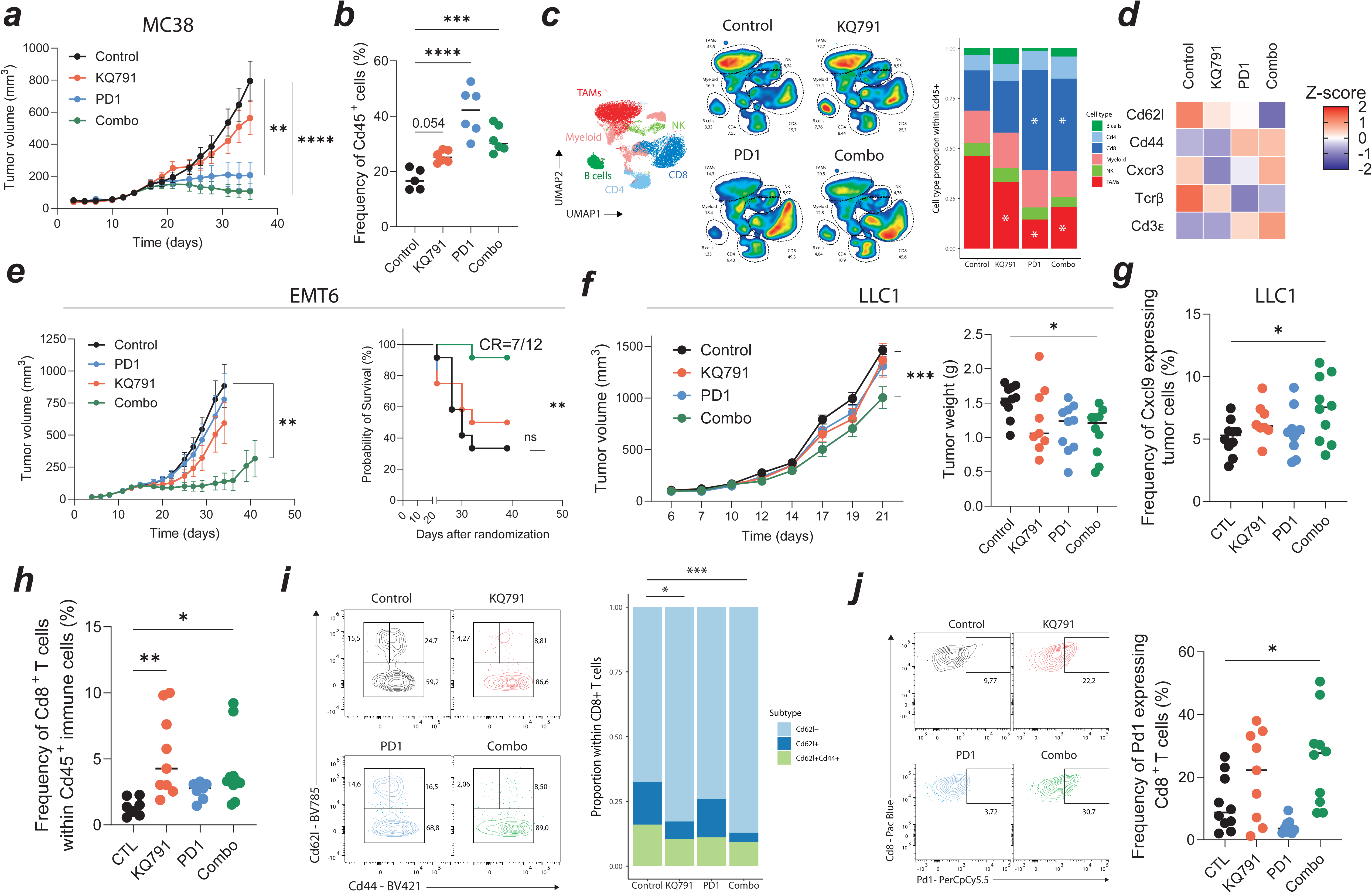
Ptpn1/2 dual inhibition sensitizes solid tumors to checkpoint blockade in syngeneic models that are either sensitive or insensitive to anti-PD1. a-d) Characterization of the tumor growth and immune infiltrates of MC38 syngeneic tumors treated with an isotype control (control), KQ791, anti-PD1, or the combination of both (Combo). a) Mean tumor growth curves of MC38 colon cancer fragments in mice treated with KQ791, anti-PD1, a combination of both (combo), or with an isotype control. Treatment regimens: KQ791 Q2DX10. anti-PD1 or control TWx3. N=10/group. b) Flow cytometric analysis of the frequency of Cd45^+^ cells in MC38 tumors. c) Flow cytometric analysis of immune populations within the Cd45^+^ compartment. Uniform Manifold Approximation and Projection (UMAP) of the major immune populations (B, Cd4^+^, Cd8^+^, Myeloid, NK cells, and Tumor-associated macrophages (TAMs))(left). Proportion graph of relative frequencies of each major immune population in MC38 tumors (right). d) Z-score heatmap of T cell markers Cd62l, Cd44, Cxcr3, and Tcrβ in tumor-infiltrating Cd8+ T cells. Gated on Cd45^+^, Tcrβ^+,^ and Cd8^+^ live cells. e) Mean growth curves of EMT6 breast cancer tumors in mice treated with KQ791, anti-PD1, a combination of both (combo), or with an isotype control. Treatment regimens: KQ791 Q2DX10. anti-PD1 or control TWx3. N=12/group (left). Kaplan-Meier curves for the number of mice reaching the tumor volume endpoint (>1000 mm^3^) (right). f-j) Characterization of the tumor growth and immune infiltrates of LLC1 syngeneic tumors. f) Mean growth curves of LLC1 lung cancer tumors in mice treated with KQ791, anti-PD1, a combination of both (combo), or with an isotype control. Treatment regimens: KQ791 Q2DX7, anti-PD1 TWx3. N=10/group (left). Dot plot of the tumor weights at day 21 (right). g) Flow cytometric analysis of Cxcl9^+^ LLC1 cancer cells (Gated on live Cd45^-^, Cd44^+^ cells). h) Frequency of Cd8^+^ T cells within the Cd45^+^ compartment as measured by flow cytometry. i) Contour plot (left) and proportion graph of the frequency of Cd62l^+^, Cd62l^-^, and Cd44^+^ cells within the Cd8^+^ T cell compartment, as measured by flow cytometry (Gated on live Cd45^+^, Tcrβ^+^, Cd8^+^ cells). j) Contour plot (left) and dot plot (right) of Pd1 expressing Cd8^+^ T cells as measured by flow cytometry (Gated on live Cd45^+^, Tcrβ^+^, Cd8^+^ cells). Error bars represent mean±s.e.m. Each dot represents an individual mouse. Statistical analysis: Tumor growth: (a,e, and f) Unpaired T-tests of the difference in tumor volume at the control endpoint. Survival analysis: (e) Log-rank (Mantel-Cox) test. Dot plots: (f,g,h and j) One-way ANOVA with Dunnett’s multiple comparison test. Key: ns nonsignificant, *p<0.05, **p<0.01, ***p<0.001 and ****p<0.0001.

We reached similar conclusions in LLC1, where only the combination of KQ791 and anti-PD1 generated a significant decrease in tumor growth (Mean growth inhibition 31%) (Fig. 5f, Extended Data Fig. 5c). Flow cytometric analyses of the LLC1 tumors revealed an enhancement of Cxcl9 by tumor cells in the Combo group (Fig. 5 g, Extended Data Fig. 5 d). In this model, we also found that the tumors treated with either KQ791 or Combo had a slight increase in the frequency of Cd8+ T cells, reaching 5.3% and 4.1% versus 1.4% in the control group, respectively (Fig. 5 g, Extended Data Fig. 5 e). In the combo group, we also found a decrease in the proportion of Cd62l+ Cd8+ T cells in the tumor infiltrates, a result in line with the previous observations made in the MC38 model (Fig. 5 i). Cd8+ T cells from the Combo group tumors also had an increased expression of PD1, indicating a potential exhaustion (Fig. 5 j). Overall, these in vivo data demonstrate that administration of Ptpn1/2 inhibitors alone is not sufficient for therapeutic responses. However, the co-administration of KQ791 with anti-PD1 sensitizes solid tumors to checkpoint blockade, even in cancer models known to be insensitive to immunotherapy. Therefore, these data also suggest that the PD1/PDL1 axis can act as a bottleneck for the efficacy of PTPN1/2 inhibitors.

### Genetic ablation of Ptpn1/2 in tumors is sufficient to generate a response to anti-PD1

Our previous results demonstrated the benefit of combining Ptpn1/2 inhibition with checkpoint blockade. To better understand the mechanism of action, however, we sought to determine whether that effect was cell intrinsic (caused by inhibition in cancer cells) or extrinsic (caused by inhibition in immune cells). To answer this question, we generated additional Ptpn1, Ptpn2, and DKO cell lines from MC38-OVA and LLC1 parental lines (Extended Data Fig. 6). Ablation of these phosphatases has a slight effect on the speed of proliferation, but the cell lines nevertheless reached confluency after 80 hours of culture (Extended Data Fig. 6 a). We also compared the three cell lines concerning their sensitivity to interferons and observed that B16-OVA, MC38-OVA, and LLC1 had the highest, intermediate, and lowest sensitivity to IFNs, respectively (Extended Data Fig. 6 b). In addition to the B16-OVA cell lines, we validated the enhancement of MHCI, Pdl1, and Cxcl9 in response to interferons in MC38-OVA and LLC1 sgCTL, sgPtpn1, sgPptn2, and sgDKO cell lines (Extended data Fig. 6 c). Given that B16-OVA was the most responsive to IFNγ, we first implanted sgCTL and sgDKO cells subcutaneously in the flank of immunocompetent mice, prior to treatment with anti-PD1 or Combo therapy (KQ791 and anti-PD1). Treatment with anti-PD1 in sgCTL B16-OVA tumors did not produce any apparent change in tumor growth (Fig. 6 a, Extended Data Fig. 7 a). On the other hand, there was a marked reduction in tumor size in sgDKO-treated tumors upon treatment with anti-PD1, indicating sensitization (Fig. 6a, Extended Data Fig. 7a). Flow cytometric analyses of the tumors at day 20 revealed an increase in immune cell infiltration (Cd45+) in sgDKO untreated and treated tumors (Fig. 6 b). Further quantification of the major immune subsets within the Cd45+ compartment revealed an increased proportion of Cd8+ T cells combined with a decreased proportion of tumor-associated macrophages (TAMs) (Fig. 6c). The Cd8+ T cells found in the TME of sgDKO tumors also had a marked increase in the levels of Cxcr3, regardless of treatment group (Fig. 6 d). Furthermore, treatment of sgDKO tumors with anti-PD1 decreased Cd62l, Pd1, and Lag3 expression in Cd8+ T cells, compared to anti-PD1-treated sgCTL tumors (Fig. 6 d). This decrease was even more pronounced in sgDKO tumors treated with the Combo therapy (Fig. 6 d). These results suggest the potential accumulation of effector-like Cd8+ T cells in Ptpn1/2 knockout tumors treated with immunotherapy. In line with this conclusion, we observed a sharp drop in the percentages of Cd62l+ Cd8+ T cells in the sgDKO tumors treated with anti-PD1 or Combo (Fig. 6 e). Given that the expression of Cxcr3 in T cells was strongly associated with sgDKO tumors, we decided to validate whether sgDKO had increased Cxcl9 production in vivo (Day 15). In line with in vitro evidence, an average of 18% of double knockout B16-OVA cancer cells expressed Cxcl9 in comparison to only 6.8% in sgCTL cells (Fig. 6 f). Given that Cxcr3 is the receptor for Cxcl9, these observations might imply that Cxcr3-positive effector T cells are recruited to tumors via a cell-intrinsic enhancement of Cxcl9 secretion. We next sought to validate those results using the LLC1 and MC38-OVA cell lines we had generated. First, we wanted to confirm that intrinsic ablation of Ptpn1/2 in tumors was sufficient for sensitization to anti-PD1. To do so, we used sgCTL or sgDKO LLC1 lung cancer tumors and treated them with PD1 checkpoint blockade. The use of LLC1 was motivated by two important characteristics: the absence of an artificial antigen (OVA) and a well-known insensibility to checkpoint blockade [51–53]. We found that only sgDKO LLC1 tumors were responsive to anti-PD1, which corroborates previous results in B16-OVA tumors (Fig. 6 g, Extended Data Fig. 7 b). We next used the sgCTL and sgDKO MC38-OVA cell lines to determine whether we would see a difference in growth and immune infiltration without treatment with anti-PD1. Unlike sgDKO B16-OVA tumors, sgDKO MC38-OVA tumors did not grow more slowly than their sgCTL counterparts (Fig. 6 h, Extended Data Fig. 7c). In accordance with previous observations, however, we did detect an increase in Cd8+ T cell infiltration and a decrease in Cd62l+ Cd8+ T cells from sgDKO MC38-OVA tumors (Fig. 6 I and j, Extended Data Fig. d and e). These results thus indicate that the deletion of Ptpn1/2 directly in tumors is sufficient for enhancing the recruitment of Cd8+ T cells in the TME. Furthermore, they highlight that the effectiveness of dual inhibitors in increasing the sensitivity of anti-PD1 is also dependent on a cancer cell intrinsic effect.

**Figure 6:**
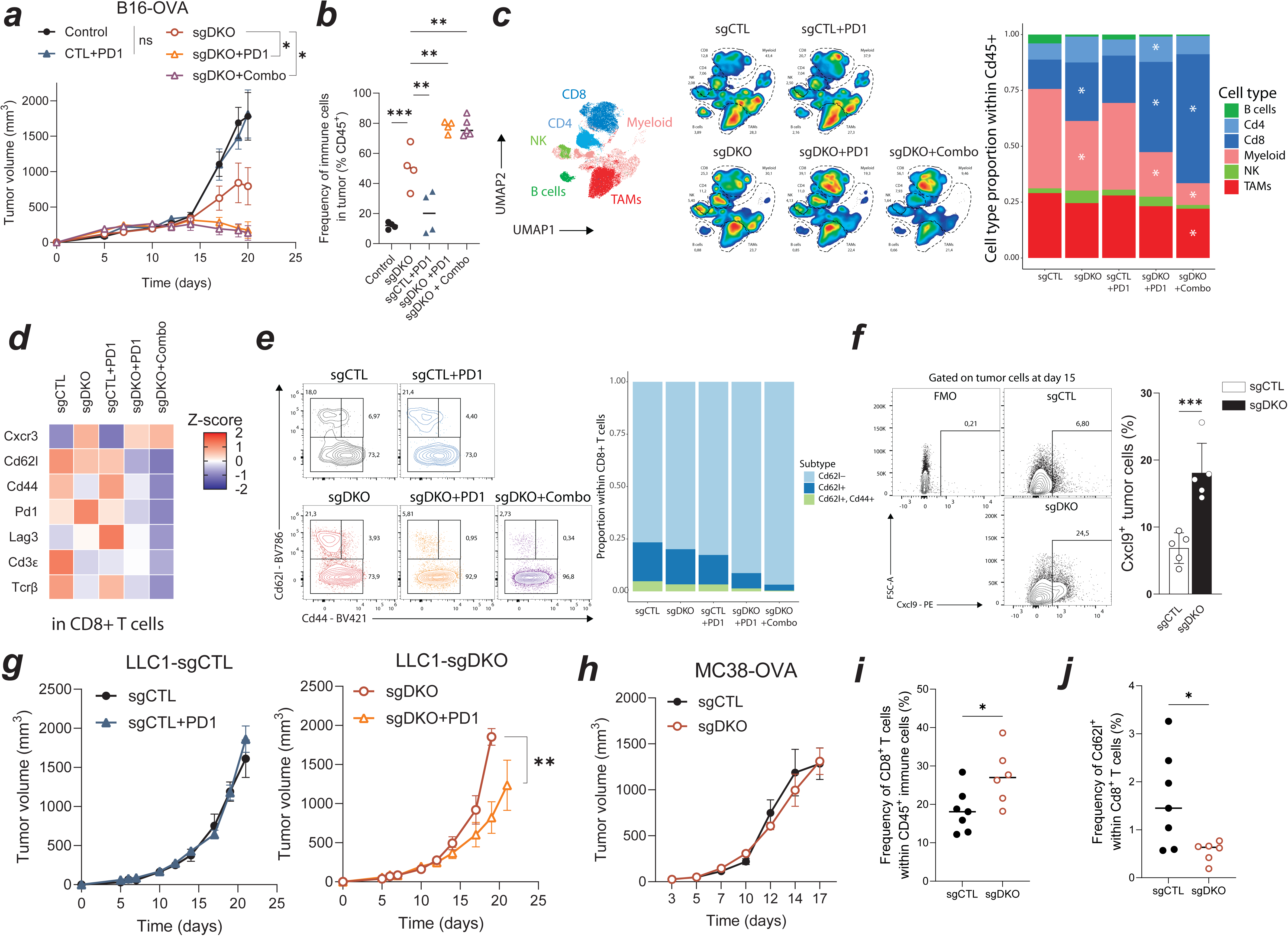
Tumor intrinsic ablation of Ptpn1/2 is sufficient for sensitization to anti-PD1. a-e) Characterization of the tumor growth and immune infiltrates of B16-OVA syngeneic tumors treated with anti-PD1, or the combination of anti-PD1 and KQ791 (Combo). a) Mean tumor growth curves of sgCTL or sgDKO B16-OVA melanomas treated with anti-PD1, anti-PD1 and KQ791 (Combo), or left untreated (control). Treatment regimens: anti-PD1 TWx2. KQ791 Q2DX6. N=4-6/group. b) Dot plot for the frequency of Cd45^+^ cells in B16-OVA tumors treated with indicated regimens as measured by flow cytometry. c) Flow cytometric analysis of immune populations within the Cd45^+^ compartment. Uniform Manifold Approximation and Projection (UMAP) of the major immune populations (B, Cd4^+^, Cd8^+^, Myeloid, NK cells, and Tumor-associated macrophages (TAMs))(left). Proportion graph of relative frequencies of each major immune population in MC38 tumors (left). d) Z-score heatmap of the T cell markers Cxcr3, Cd62l, Cd44, Pd1, Lag3, Cd3ε, and Tcrβ in tumor-infiltrating Cd8+ T cells. Gated on Cd45^+^, Tcrβ^+,^ and Cd8^+^ live cells. e) Contour plot (left) and proportion graph of the frequency of Cd62l^+^, Cd62l^-^, and Cd44^+^ cells within the Cd8^+^ T cell compartment as measured by flow cytometry (gated on live Cd45^+^, Tcrβ^+^, Cd8^+^ cells). f) Contour plot (left) and dot plot (right) of Cxcl9^+^ B16-OVA cells 15 days after tumor inoculation as measured by flow cytometry (gated on live Cd45^-^ tumor cells). g) Mean growth curves of sgCTL or sgDKO LLC1 lung cancer tumors treated with or without anti-PD1. Treatment regimens: anti-PD1 TWx2. N=4-6/group. h) Mean growth curves of sgCTL or sgDKO MC38-OVA tumors (no treatment). N=6/group. Error bars represent mean±s.e.m. Each dot represents an individual mouse. Statistical analysis: Tumor growth: Unpaired T-tests of the difference in tumor volume at the last time point (a,g, and h). Dot plots: One-way ANOVA with Dunnett’s multiple comparison test (b). Unpaired T-tests (f, I, and j). Key: ns nonsignificant, *p<0.05, **p<0.01, ***p<0.001 and ****p<0.0001.

### Ptpn1/2 dual deficiency sensitizes cancer cells to IFNγ & TNFα-mediated cell death

Cd8+ T cells induce cytolysis of their targets upon direct recognition, but their activation also leads to the production of IFNγ & TNFα, which are important for ensuring the apoptosis of their target cells. In our flow cytometric analysis of tumor-infiltrating Cd8+ T cells, we found an enhancement of IFNγ & TNFα expression in T cells from sgDKO B16-OVA tumors compared to controls (23% of Cd8^+^ were IFNγ^+^ TNFα^+^ in sgCTL versus 33% in sgDKO tumors) (Fig. 7 a). This result suggests that T cells encountering Ptpn1/2 null tumors are activated more robustly than their control counterparts. This result also prompted us to explore whether Ptpn1/2 deficient cancer cells were also intrinsically more susceptible to IFNγ & TNFα-mediated cell death. To do so, we first cultured B16-OVA cells of the various Ptpn1/2 genotypes with exogenous IFNβ or IFNγ, with or without TNFα (Fig. 7 b). We quantified cell abundance after 48-72h of cell culture using a fluorescence-based method (Cyquant). After 48 hours, we only noticed a drop in cell abundance in response to IFNγ in sgPtpn1, sgPtpn2, and sgDKO lines (Fig. 7 b). The decrease in cell abundance was most drastic when cells were co-treated with TNFα, especially in sgDKO cultures, where the cell abundance was less than 25% of what it was in untreated cells (Fig. 7 b). sgDKO started to exhibit morphological differences, such as cell rounding and swelling, 24 hours post-IFNγ/TNFα treatment (Extended Data Fig. 8 a). When culturing the cells for up to 72h, we observed a relative cell abundance of 15% and 5% in sgCTL and sgDKO cells treated with IFNγ/TNFα, respectively (Extended Data Fig. 8 b). These results thus indicated that Ptpn1/2 deficient cells responded to IFNγ/TNFα treatment faster than the control cells. To quantify the amount of cell death resulting from IFNγ/TNFα treatment, we next proceed to use Lactate Dehydrogenase (LDH) release assays. After 48 hours of IFNγ/TNFα treatment, we found an additive effect between Ptpn1/2 deletion and LDH release (Fig. 7 c). Indeed, while single knockout cells had a slight increase of 20-30% in LDH release in response to IFNγ/TNFα treatment, dual deletion led to a more than 50% increase (Fig. 7 c). We also found an enhancement of LDH release in sgDKO LLC1 lung cancer cells compared to controls, thus validating the presence of this phenotype in different cancer types (Extended Data Fig. 8 c). Pharmacological inhibition assays revealed the dependence of the activity of JAK1-3, but not SFKs, for this phenotype (Extended Data Fig. 8 d). Cell death was also accompanied by an increase of iNos, Gasdermin D (Gsdmd), and Caspase 11 in sgDKO cells (Extended Data Fig. 8 e). Most strikingly, we found that treatment with IFNγ was sufficient for inducing Caspase 11 expression in sgPtpn2 or sgDKO cells, while both IFNγ & TNFα were required in sgCTL or sgPtpn1 cells (Extended Data Fig. 8 e). Overall, these results demonstrate that deletion of Ptpn1/2 enhances the sensitivity of cancer cells to IFNγ/TNFα-mediated cell death in a gene-dose-dependent manner.

**Figure 7:**
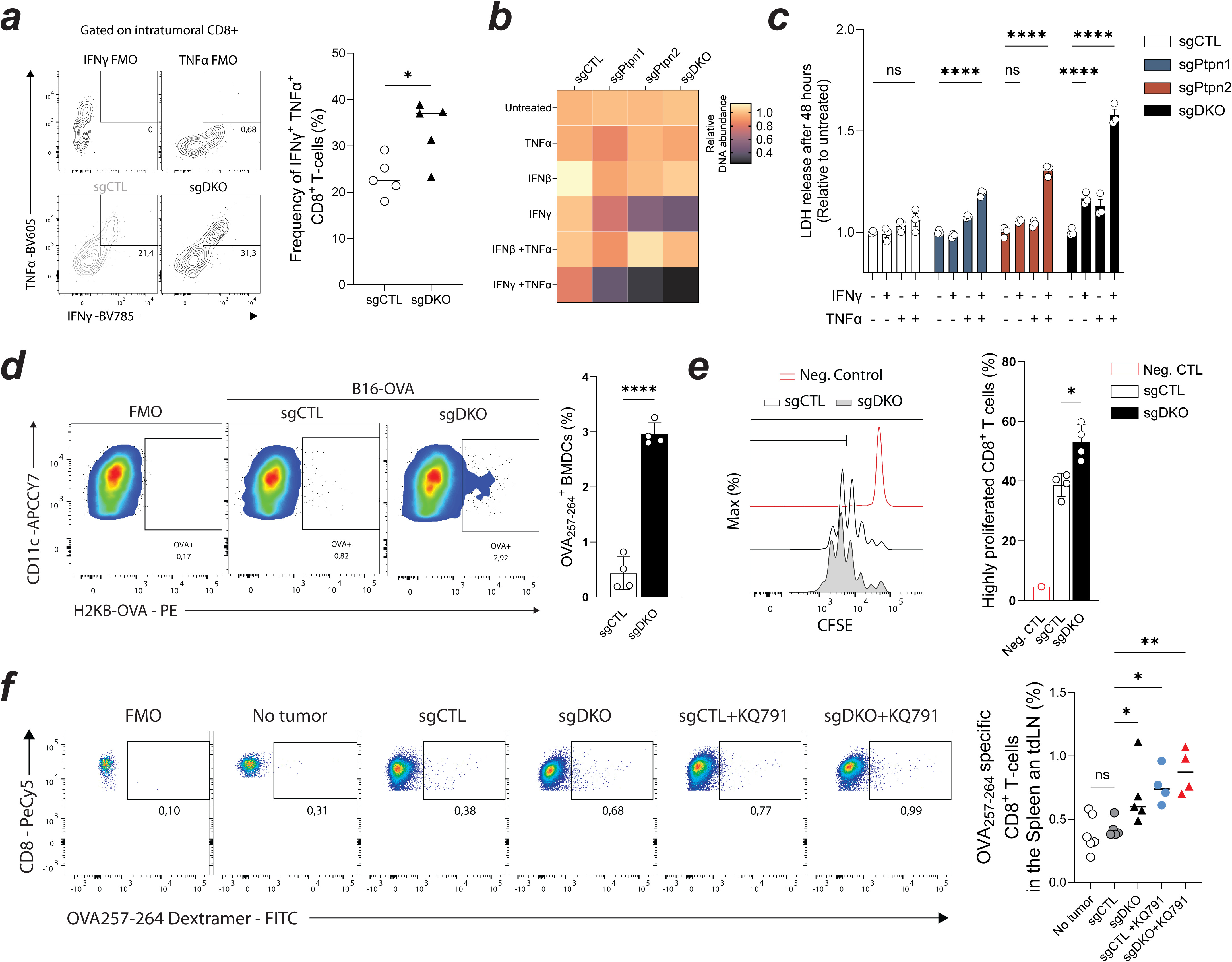
Ptpn1/2 hinder antigenic release by cancer cells and prevent the establishment of cellular immunity. a) Contour plot (left) and dot plot (right) of IFNγ and TNFα positive Cd8^+^ T cells 15 days after tumor inoculation as measured by flow cytometry (gated on live Tcrβ^+^, Cd8^+^ cells). b) Heatmap of the relative DNA abundance of sgCTL, sgPtpn1, sgPtpn2, and sgDKO B16-OVA cells cultured with IFNβ, IFNγ, and/or TNFα for 48h. Data was normalized to the untreated cells in each genotype. c) Relative LDH release in sgCTL, sgPtpn1, sgPtpn2, and sgDKO B16-OVA cultures with IFNγ, and/or TNFα (48h post-stimulation). Data was normalized to the untreated cells in each genotype. d) Flow cytometric analysis of OVA_257-264_ positivity in bone marrow-derived dendritic cells (BMDCs) cocultured with sgCTL of sgDKO B16-OVA cells (left) and associated bar graph (right). e) CFSE proliferation histograms of OT-1 T cells co-cultured with BMDCs harvested from sgCTL of sgDKO B16-OVA co-cultures (left). Bar graph of the frequency of highly proliferated Cd8^+^ T cells from the CFSE analysis (CFSE MFI<5x10^3^) (right). f) Flow cytometric analysis of OVA257-264 dextramer positivity in CD8+ T cells from spleen and tumor-draining lymph node (tdLN) of mice inoculated with sgCTL or sgDKO tumors and treated with or without KQ791 (day 15). Data representative of at least three independent experiments (b-e). Error bars represent mean±s.e.m. Each dot represents an individual mouse (a and f). Statistical analysis: Unpaired T-tests (a,d, and e). Two-way ANOVA with Tukey’s multiple comparisons (b and c). One-way ANOVA with Dunnett’s multiple comparison test (f). Key: ns nonsignificant, *p<0.05, **p<0.01, ***p<0.001 and ****p<0.0001.

### Ptpn1/2 hinder antigenic release by cancer cells and prevent the establishment of cellular immunity

We next hypothesized that the death of sgDKO cells could also be relevant for antigen release. To test this hypothesis, we cultured sgCTL or sgDKO B16-OVA cells with mature bone marrow-derived dendritic cells (BMDCs) and quantified the abundance of the OVA-derived peptide (257-265) in complex with MHCI molecules of the BMDC’s surface (Fig. 7d). We found that around 3% of BMDCs cultured with sgDKO cells for 24 hours were positive for OVA_257-264_, while observing only 0.43% positivity in sgCTL counterparts (Fig. 7d). These BMDCs were next harvested and used to prime OT-1 (OVA_257-264_ restricted Cd8+ T cells) cells. As expected, OT-1 cells that were primed with the BMDCs previously co-cultured with sgDKO cancer cells proliferated more than the sgCTL fed counterparts (Fig. 7e). These results thus suggest that Ptpn1/2 deficient cancer cells either release more antigens or are processed more easily by antigen-presenting cells, enhancing antigen-specific T cell responses. In previous in vivo experiments, we had also noticed an increase in the frequency of OVA_257-264_ positive dendritic cells in mice that were bearing sgDKO B16-OVA tumors (Extended Data Fig. 9a). Still, it remained unknown whether KQ791 could also enhance the antigen cross-presentation capabilities of DCs. Our group has previously identified that Ptpn1/2 suppresses dendritic cell differentiation and maturation via STAT1 and STAT4 dephosphorylation [11]. To quantify antigen cross-presentation, we used BMDCs cultures pulsed directly with the OVA_257-264_ peptide. Treatment with KQ791 significantly increases the frequency of OVA_257-264_-MHCI complexed on the surface of the BMDCs (Extended Data Fig. 9 b). Ptpn1/2 inhibition also enhanced the expression of MHC2, Cd86, and Pdl1 on the surface of BMDCs (Extended Data Fig. 9c). In human monocyte-derived dendritic cells, PTPN1/2 inhibition enhanced the secretion of TNFα, IL10, and CXCL9 in response to monophosphoryl lipid A (MPLA) (Extended Data Fig. 9d). Priming of OT-1 cells with KQ791-treated BMDCs substantially enhanced their proliferation and their expression of the activation marker Cd25 (Extended Data Fig. 9e and f). This evidence demonstrates that Ptpn1/2 inhibition also enhances the functionality of antigen-presenting cells, which could be an underlying determinant of the response to Ptpn1/2 inhibitors. Finally, we sought to validate these two sets of findings (increased antigen release and DCs activity in response to Ptpn1/2 inhibition) in vivo using B16-OVA tumors. To do so, we inoculated mice with sgCTL or sgDKO tumors and treated the animals either with or without KQ791 (100mg/kg every two days for two weeks). At 15 days, which represents the peak of adaptive immune responses, we harvested the spleen and tumor draining lymph nodes (tdLNs) of these animals and quantified the frequencies of OVA_257-264_ specific Cd8+ T cells (Fig. 7f). We found that sgDKO tumor-bearing mice had a significant increase in the frequency of peripheral OVA_257-264_ specific Cd8+ T cells (Fig. 7f). Treatment of sgCTL tumors with KQ791 yielded a similar effect, almost doubling the frequency of OVA_257-264_ specific Cd8+ T cells compared to sgCTL tumor-bearing untreated animals (Fig. 7f). Finaly, treatment of sgDKO tumor-bearing animals with KQ791 yielded the strongest increase in OVA_257-264_ specific Cd8+ T cells, potentially by combining increased antigen release with enhanced DC activity (Fig. 7 f). The collection of these data thus suggests that Ptpn1/2 normally hinders the immunogenicity of cancer cells, whereas Ptpn1/2 ablation or inhibition increases the targetability of cancer cells and improves the generation of anti-tumor immunity.

## Discussion

Our study revealed that the Ptpn1 and Ptpn2 phosphatases directly contribute to cancer cell immune evasion in established tumors. By decreasing the signaling of Type I and Type II interferons, PTPN1/2 limit the abundance of MHC-class I molecules on the surface of cancer cells, thus impeding their recognition by Cytotoxic T lymphocytes. On the other hand, dual ablation or inhibition of these phosphatases also raises the expression of the major checkpoint receptor ligand PDL1 on the surface of cancer cells, representing an important limitation for these targets. This increased PDL1 expression is mediated by STAT1/3 signaling and is likely responsible for the limited efficacy of KQ791 monotherapy. Combination with anti-PD1 or anti-PDL1 is thus required for maximal efficacy, since these checkpoints could induce resistance to PTPN1/2 inhibition [54]. Other small-molecule inhibitors, such as ABBV-CLS-484, have achieved monotherapeutic effects in some cancer models [10, 13]. ABBV-CLS-484 is different from KQ791 in many regards, such as solubility, IC_50,_ and inhibition of PTPN9. Most notably, KQ791 is 10 times more selective to PTPN1/2 over PTPN9 compared to ABBV-CLS-484 [10]. Small-molecule degraders of PTPN1/2 also affect PTPN9 more than any other phosphatase, which highlights a common challenge when targeting the catalytic site of PTPN2 [55]. More investigations are required for understanding the relevance of PTPN9 for cancer immunotherapy and whether it is a desirable target when inhibiting PTPN1/2. PTPN1/2 and PTPN9 have some common functions, such as STAT3 dephosphorylation [56], HER2/ErbB2 signaling [57] and iron metabolism [58, 59]. Additional targeting of PTPN9 might also lead to more severe adverse effects since homozygous knockout in mice (Meg2−/−) leads to embryonic lethality and defects in T cell and platelet activation [60].

Besides enhancing MHC-class I expression on cancer cells, PTPN1/2 inhibition greatly enhanced CXCL9 secretion in response to IFNγ. CXCL9 and CXCL10 are primarily produced by macrophages in the TME and are required for the efficacy of immune checkpoint blockade [61, 62]. CD8+ T cell recruitment to tumors depends on CXCL9/10 secretion [61]. The CXCR3-CXCL9/10 axis also stimulates the contact between antigen-specific CD8+ T cells and dendritic cells within tumors, leading to efficacious anti-cancer T cell immunity [62]. In our study, we have demonstrated that the production of CXCL9 can also be artificially induced by tumor cells and dendritic cells using genetic deletion or inhibition of PTPN1/2. The enhancement of CXCL9 production might have led to the accumulation of CXCR3+ effector T cells, which in turn increases the proportion of active CTLs within the TME. Our group and others have also demonstrated that the inhibition of PTPN1/2 in T cells enhances their proliferation and limits their functional exhaustion [5, 12, 13, 63]. Therefore, PTPN1/2 inhibition provides multiple benefits to cancer immunotherapy by acting simultaneously on the immune system and in cancer cells. However, we have found that the PD1 axis still acts as a bottleneck during PTPN1/2 inhibition (Fig. 5). Furthermore, Ptpn1/2 ablation in tumors was sufficient for therapeutic sensitization to anti-PD1 (Fig. 6).

We also observed that ablation of PTPN1/2 in cancer cells is sufficient for enhancing antigen presentation by dendritic cells, and potentially other antigen-presenting cells. Genetic deletion or pharmacological inhibition of these phosphatases increases the frequency of antigen-specific T cells in the periphery (Fig. 7). Although the exact molecular mechanisms for this enhancement of the immunogenic cell death pathway remain to be elucidated, we have found that the abolition of PTPN1/2 activity enhances the expression of several core components of the TNFR1 signalling cascade. Identified DEGs in sgDKO cells, such as *Ripk1*, *Casp8*, *Casp11*, *Gsdmd*, and *Nos2*, could be linked to the enhancement of immunogenic cell death by promoting necrosis and pyroptosis rather than apoptosis in response to IFNγ/TNFα stimulation. It has been previously identified that STAT1 and IRF1 orchestrate the transcriptional response necessary for IFNγ/TNFα-induced PANoptosis [64, 65]. On the other hand, activation of the RIPK1/FADD/CASP8 axis by nitric oxide produced by iNOS (NOS2) was shown to be the crucial event leading to PANoptosis downstream of STAT1 and IRF1 [64]. PTPN1/2 ablation or inhibition thus increases the expression of core components responsible for inducing PANoptosis, such as *Ripk1*, *Casp8*, and *Nos2*, which in turn enhances the susceptibility of cancer cells to IFNγ and TNFα.

**In conclusion**, we have demonstrated the immunosuppressive roles of PTPN1/2 in cancer cells. Inhibition of PTPN1/2 in cancer cells is sufficient for sensitizing different types of solid tumors to checkpoint blockade immunotherapy. While PD1 alleviates the exhaustion of antigen-experienced T cells, PTPN1/2 inhibition allows for the generation of effector T cells and their recruitment to tumors, most likely through a Cxcl9-dependent manner. Therefore, the effects of the two approaches are complementary to one another and generate potent immune responses against solid tumors. The use of small-molecule PTPN1/2 inhibitors could thus increase the number of patients potentially benefiting from ICB and expand its use to new cancer types.

## Material & Methods

### Mice and cell lines

All mouse procedures were performed using 8-15 weeks-old mice in accordance with the Canadian Council on Animal Care ethical regulations and the McGill University Research and Ethics Animal Committee. C57BL/6J (Jackson Laboratory, 000664) and C57BL/6-Tg (TcraTcrb)1100Mjb/J (OT-1) (Jackson Laboratory, 003831) mice were housed and bred in the Comparative Medicine and Animal Resources Centre (CMARC). The B16-F10 expressing chicken ovalbumin (OVA) murine melanoma cell line, B16-MO4, was purchased from Millipore Sigma (SCC420). MC-38, LLC1, HEK293T/17, MiaPaca2, MCF-7, OVCAR-3, U251, U-87-MG, A549, HCT116, Hela 299, A431 and LnCap cell lines were purchased from ATCC. All cell lines, except OVCAR-3 cells, were grown in DMEM high glucose (Cytiva HyClone, SH30022FS), 10%FBS, 1XPenicillin Streptomycin (Cytiva Hyclone, SV30010). OVCAR-3 cells were grown in RPMI (Cytiva HyClone, SH3002701), 20% FBS, 1XPenicillin Streptomycin, and 1X Insulin-Transferrin-Selenium (Gibco, 41400045). Cell lines were tested for mycoplasma prior to culture and repeatedly using PCR.

### DiFMUP (6,8-Difluoro-4-Methylumbelliferyl Phosphate) assays for phosphatase activity

Enzyme reactions were conducted in assay buffer 50mM HEPES pH7.0, 3mM DTT, and 1mg/mL BSA using the purified recombinant GST-PTP1B and GST-TC-PTP catalytic domains [66]. DiFMUP was used as a substrate. Determination of kinetics constants using DiFMUP as substrate: The hydrolysis of DiFMUP was conducted in black 96-well plates (Corning) in a final volume of 100µL at 25°C. The reaction was monitored by measuring the fluorescence (excitation wavelength 358nm/emission 455nm) with the Varioskan plate reader (Thermo Electron). Enzyme dilution was determined by choosing a reaction rate to comprise in an Absorbance range of 5-30 FU units/min. For the kinetic assays, fluorescence was monitored over 10 minutes in 30-second intervals, and rates were calculated using a non-linear least-square fitting procedure. Km was determined from rates at various substrate concentrations using the Michaelis-Menten equation and IC50 values were derived by a sigmoidal dose-response (variable slope) curve using GraphPad Prism software. A substrate concentration equivalent to the Km value was used for IC50 determinations. KQ-791 and L598 Inhibitors were diluted in PBS, and then a serial dilution starting from 24µM and covering 3 log scales was made in assay buffer.

### Gene silencing and CRISPR-mediated knockout

For knocking down Ptpn1 and Ptpn2 in mouse B16-MO4 cells, we transfected the equivalent of 25 pmol of siRNA with the equivalent volume of Lipofectamine RNAiMAX reagent (Thermo Fisher Scientific, 13778150) for 1x10^5^ cells in 24 well plates. The setup was scaled proportionally for larger plates. We used siGENOME Mouse *Ptpn1* (Dharmacon, M-040818-01-0005) and *Ptpn2* (Dharmacon, M-040061-01-0005) smartpool siRNAs for the knockdown of the phosphatases and siGENOME Non-Targeting Control siRNAs, #1 (Dharmacon, D-001210-01-05) as the negative control.

For genetic knockout, we used a bulk CRISPR/Cas9 approach using the lentiCRISPRv2 all-in-one expression vector (Addgene, 52961). LentiCRISPR v2 was a gift from Feng Zhang (Addgene plasmid # 52961 ; http://n2t.net/addgene:52961; RRID:Addgene_52961)[67]. RNA primers were cloned into the lentiCRISPRv2 backbone using a Golden Gate reaction as per [68]:

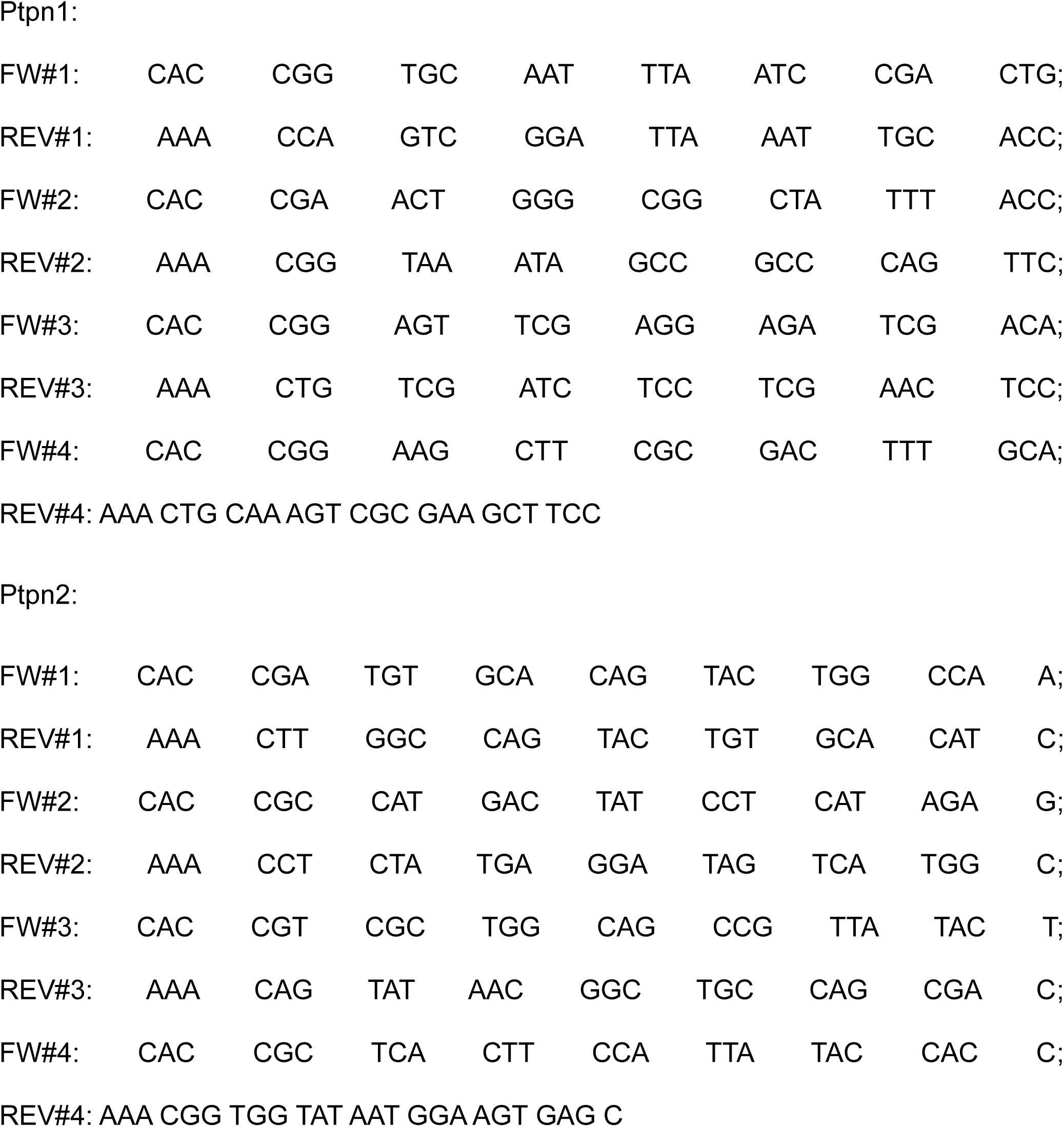

1uL of each forward and reverse oligonucleotide was annealed using the T4 polynucleotide kinase (New England Biolabs, M0201L) at 37 C for 30min, 95 C for 5 minutes, followed by a cooling down to 25 C at a rate of 5 C/min. After cloning, the plasmids were validated by pyrosequencing (Génome Québec). Lentiviral particles were prepared using transfection of PAX2, VSV-G, and lentiCRISPRv2 with the Ptpn1/2 gRNAs in HEK293T/17 cells. The supernatant was collected every day for 48 hours. Supernatants were then filtered through a Filtropur 0.45 μm syringe filter (Sarstedt, 83.1826). Viral supernatants were then concentrated using Viro-PEG lentivirus concentrator (OZBiosciences, LVG100) as per the manufacturer’s instructions. Cells were then transduced with a high multiplicity of infection (MOI) using 8 μg/mL of polybrene (Millipore Sigma, TR-1003). Infected cells were incubated for 48-72h before selection with 5 μg/mL puromycin (BioShop Canada, PUR333.25) for an additional 72 hours. We chose to use set#4 and set#1 for Pptn1 and Ptpn2, respectively, based on knockout efficiency in the DKO cells. Both of these guide RNAs target exon 1 of the respective phosphatase in the Mus Musculus genome. We used lentiCRISPRV2 cloned with a sgControl (CACCgGCACTACCAGAGCTAACTCA) as the transduction for the control cell lines sgCTL (Addgene, 125836). lentiCRISPR v2-sgControl was a gift from Boyi Gan (Addgene plasmid # 125836 ; http://n2t.net/addgene:125836; RRID:Addgene_125836) [69].

### Cytokine/chemokine profiling

B16-MO4 cells were silenced for Ptpn1/2 as previously stated and were seeded at 1x10^6^ cells per well of a 6-well plate. After the knockdown period (48 hours) cells were stimulated with 20ng/mL of recombinant murine IFNγ (Preprotech, 315-05) for 24 hours. Supernatants were collected, and cytokine/chemokine abundance was measured with the Proteome Profiler Mouse Cytokine Array Kit, Panel A (R&D, ARY006), as per the manufacturer’s instructions.

### Cell stimulations

B16, LLC1 or MC38 cells were initially seeded at 2x10^5^ cells/mL, left to adhere for 24-48h prior to stimulation with 20ng/mL of recombinant murine IFNβ1 (BioLegend, 581302), IFNγ (Preprotech, 315-05) for 24 hours (Flow cytometry) or 30 minutes (Western Blotting). Human cancer cells were seeded at 1x10^5^ cells/mL for 48 hours prior to stimulation with 20ng/mL of recombinant Human IFN-γ (Preprotech, 300-02) for 24 hours prior to flow cytometric analysis.

### Flow cytometry

For flow cytometric analysis, 2x10^5^ to 5x10^5^ cells were harvested for processing. Adherent cells were detached using an Enzyme-free cell dissociation buffer (Life Technologies, 13151014). Staining and washing were performed in round bottom polystyrene 5mL tubes (Falcon, 14-959-6). Cells were washed once using PBS prior to life-death staining with the Fixable Viability Dye eFluor™ 780 (Biosciences, 65-0865-14). Cells were stained for 20 minutes on ice prior to a single wash with flow cytometry staining buffer (PBS, 2% heat-inactivated FBS, 500μM EDTA, and 0.02% m/v Sodium Azide). When staining immune cells, the CD16/32 Fc receptors were blocked for 15 minutes using a 1 in 300 dilution of Anti-Mouse CD16/CD32 (Fc block) (Life Technologies, 14-0161-85). Surface staining was performed on ice for 45 minutes in flow cytometry staining buffer. A specific dilution of fluorochrome-conjugated antibodies was used for each antibody (See list of antibodies in the antibodies section). After surface staining, samples were washed once with flow cytometry staining buffer and resuspended in 200uL of flow cytometry staining buffer prior to analysis. Intracellular staining was realized using the Bd cytofix/cytoperm Fixation/Permeabilization Solution Kit (BD, 554714) as per the manufacturer’s instructions. Samples were acquired with a 4 or 5-laser LSR Fortessa or with a Cytek Aurora Spectral Flow Cytometer and analysed using FlowJo V10.10. The *Downsample* and *UMAP* plugins were used for dimensionality reduction analysis on the immune infiltrate in tumors (Cd45^+^ cells).

For the staining of OVA_257-264_ positive T cells, we stained 1 × 10^6^ cells from the spleen and lymph nodes of tumor-bearing animals using an OVA-MHC dextramer (Immudex, JD02163, FITC). Staining was performed as per the manufacturer’s protocol (CD3/CD8 pre-staining followed by dextramer staining at room temperature). Reagents and staining conditions were validated using OT-1 T cells as a positive control.

Antibodies used (Supplier, Catalog#, Dilution factor):

|  |  |
| --- | --- |
| Cd19-AF647 | (BioLegend,115525,1:1600) |
| B220-AF700 | (BioLegend,103232,1:800) |
| NK1.1-BV605 | (BioLegend,108739,1:200) |
| Pd-1-PerCPCy5.5 | (BioLegend,109119,1:200) |
| Cd3-APC/Fire810 | (BioLegend,100267,1:150) |
| Cd4-PE/Dazzle 594 | (BioLegend,100455,1:800) |
| Cd8a-Pacific Blue | (BioLegend,100728,1:600) |
| Cd44-BV421 | (BD,563970,1:400) |
| Cd62L-BV786 | (BD,564109,1:1200) |
| Cd45-FITC | (BioLegend,147709,1:300) |
| Tcrb-APC | (BioLegend,109212,1:200) |
| Ter119-BV650 | (BioLegend,116235,1:400) |
| Cxcr3-PE | (BioLegend,155903,1:200) |
| Ly6G-PE/Cy7 | (BioLegend,127617,1:400) |
| Ly6C-APC/Cy7 | (BD,560596,1:400) |
| CD11b-SparkYG593 | (BioLegend,101281,1:800) |
| CD11c-BV510 | (BioLegend,117337,1:150) |
| F4/80-PE/Cy5 | (BioLegend,123111,1:600) |
| MHCII-Spark Blue 550 | (BioLegend,107661,1:800) |
| H-2Kb (MHCI)-FITC | (BioLegend,116505,1:300) |
| H-2Kb (MHCI)-PerCPCy5.5 | (BioLegend,116515,1:300) |
| H-2Kb (MHCI)-AlexaFluor 700 | (BioLegend,116521,1:300) |
| H-2Kb bound to SIINFEKL-PE | (BioLegend,141603,1:300) |
| H-2Kb bound to SIINFEKL-APCCy7 | (BioLegend,141607,1:300) |
| anti-mouse Pdl1 (Cd274)-APC | (BioLegend,124311,1:300) |
| anti-mouse Cxcl9-PE | (BioLegend,515603,1:200) |
| HLA-A,B,C-AlexaFluor 700 | (BioLegend,311437,1:300) |
| anti-human PDL1 (CD274)-BV605 | (BioLegend,329723 ,1:300) |
| PE Mouse Anti-Human MIG (CXCL9)-PE | (BD,566013,1:200) |
| APC anti-human CXCL10 (IP-10)-APC | (BD,519505,1:100) |
| APC anti-human CXCL9 (MIG)-APC | (BioLegend,357906,1:200) |
| Cd11c-APCCy7 | (BioLegend,117323,1:300) |
| anti-mouse Tnf-α-BV605 | (BioLegend,506329,1:100) |
| anti-mouse IFN-γ -BV785 | (BioLegend,505838,1:100) |
| Cd25-PE | (BioLegend,101904,1:400) |
| Cd8a-PeCy5 | (BioLegend,100709,1:600) |
| Cd3e-FITC | (BioLegend,152303,1:300) |

### In-vitro OT-1 cytotoxicity assays

To obtain CD8^+^ OT-1 T cells, spleens of OT-1 mice were macerated on 70μm strainers (Falcon, 352350) in PBS 2% FBS prior to centrifugation at 300g for 5 minutes. Red blood cell lysis was then performed on the cell pellet using 2mL of ACK lysis buffer (Gibco, A1049201) as per the manufacturer’s guidelines. OT-1 splenocytes were then expanded using 1ug/mL OVA_257-264_ peptide (SIINFEKL, Sigma-Aldrich, S7951) at a density of 2.5 x 10^5^ cells. After 48 hours, blasts were expanded with 100 U/mL of recombinant murine IL-2 (Preprotech, 212-12) in fresh media and harvested on day four. Two days prior to co-culture with OT-1 cells, B16-MO4 cells stained with CellTrace™ CFSE Cell Proliferation Kit (Life Technologies, C34554) and seeded at 5 x 10^5^ cells per well of a 6-well plate, with or without 20μM of KQ791.

### Protein Aggregation Capture (PAC) of Proteins with On-Bead Digestion

Samples were first reduced with 9 mM dithiothreitol in 50mM ammonium bicarbonate incubated on an Eppendorf ThermoMixer, 37°C for 30 minutes, 700 RPM and, after cooling for 5 minutes, alkylated with 17 mM iodoacetamide in 50mM ammonium bicarbonate at room temperature for 30 minutes, 700 RPM in the dark. Residual nucleic acids were digested using 800 units of benzonase per sample in the presence of 2 mM MgCl₂, incubated at 37°C for 90 minutes at 700 RPM. Next, Protein Aggregation Capture with Hydroxyl beads from ReSyn Biosciences was performed using 125μl of washed beads per sample. After that, a volume of 100% acetonitrile was added to obtain a 50% final concentration to initiate protein binding. The samples were incubated for 30 minutes, at room temperature, 1000 RPM and placed on a magnetic rack for 2 minutes. The supernatants were discarded and kept for further analysis. The beads were washed three times with 70% ethanol. After the last supernatant was discarded, the beads were resuspended with 500μl of Lys-C/Trypsin Mix from Promega (0.016ug/μl in 50mM ammonium bicarbonate). The samples were sonicated in a water bath for 5 minutes and digested overnight at 37°C, 700 RPM. After the digestion is completed, the samples were placed on a magnetic rack and the supernatants were transferred to Eppendorf LoBind tubes. The beads were rinsed with 200μl of 50 mM Ammonium bicarbonate and the supernatants were pooled.

### LC-MS/MS analysis

Samples were resolubilized under agitation for 15 min in 12 µL of 1%ACN / 1% formic acid and loaded onto a 75 μm i.d. × 150 mm Self-Pack C18 column installed in the Easy-nLC II system (Proxeon Biosystems). The buffers used for chromatography were 0.2% formic acid (buffer A) and 100% acetonitrile/0.2% formic acid (buffer B). Peptides were eluted with a two-slope gradient at a flow rate of 250 nL/min. Solvent B first increased from 1 to 33% in 125 min and then from 33 to 70% in 9 min. The HPLC system was coupled to Orbitrap Fusion mass spectrometer (Thermo Scientific) through a Nanospray Flex Ion Source. Nanospray and S-lens voltages were set to 1.3-1.8 kV and 60 V, respectively. Capillary temperature was set to 250 °C. Full scan MS survey spectra (m/z 350-1500) in profile mode were acquired in the Orbitrap with a resolution of 120,000 and a target value at 4e5. The most intense peptide ions were fragmented in the HCD cell with a normalized collision energy of 30%, and the fragment ions were analyzed in the linear ion trap with a target value at 1e4. The duty cycle was set to 3 seconds, and target ions selected for fragmentation were dynamically excluded for 9 sec. An MS3 scanning was performed upon detection of a neutral loss of phosphoric acid (48.99, 32.66 or 24.5 Th) in MS2 scans. Protein database searching was performed with Mascot 2.6 (Matrix Science) against the Uniprot Mus Musculus protein database (version March 1st, 2023). The peak list files were generated with Proteome Discoverer (version 2.3) using the following parameters: minimum mass set to 500 Da, maximum mass set to 6000 Da, no grouping of MS/MS spectra, precursor charge set to auto, and minimum number of fragment ions set to 5. The mass tolerances for precursor and fragment ions were set to 10 ppm and 0.6 Da, respectively. Trypsin was used as the enzyme, allowing for up to 3 missed cleavages. Cysteine carbamidomethylation was specified as a fixed modification, and methionine oxidation, serine, threonine and tyrosine phosphorylation modifications as variable modifications. Data analysis was performed using Scaffold (version 4.8).

### Bulk mRNA sequencing and data processing

Two days prior to RNA extraction, 1x10^6^ B16-MO4 cells per condition (CTL, sgPtpn1, sgPtpn2, sgDKO, and 20μM KQ791) were seeded in triplicates in 6-well plates. Cells were left to attach overnight before stimulation with 20ng/mL of recombinant murine IFNβ1 (Biolegend, 581302) for 6 hours. Supernatants were removed, and plates were kept frozen at -80C until RNA extraction. mRNA isolation was realized with RNAEasy plus mini kit (Qiagen, 74134) as per the manufacturer’s instructions. RNA yield was quantified with a Thermo Scientific NanoDrop 1000. 1000ng of RNA was sent to the Centre d’expertise et de services Génome Québec for quality control, library preparation, and sequencing. Libraries were prepared using the Illumina TruSeq RNA Library Prep Kit following the manufacturer’s instructions. High throughput sequencing was performed using an Illumina NovaSeq PE100. The NEBNext® Multiplex Oligos for Illumina® (Dual Index Primers i7 and i5) were used as adaptors for Illumina sequencing. 28 to 46 million reads were obtained per sample, averaging 35 million reads. The resulting reads were aligned and quantified to the GRCm38 mouse reference genome assembly using Salmon [70].

Analysis: Raw counts were normalized to transcripts per million using *tximport* and analyzed for differential gene expression using DESeq2 [71, 72]. Nominal p-values were corrected for multiple testing using the Benjamini-Hochberg False Discovery Rate method (FDR). Gene ontology enrichment analysis was performed using the ClueGO app in Cytoscape. Volcano/dot plots and heatmaps were created using the R packages *ggplot2* and *Complex Heatmap*, respectively.

### Western blotting

Cells were washed using PBS and kept at -80C until lysis. Cells were lysed in RIPA buffer (1% Triton X-100, 150 mM NaCl, 1 mM NaVO_4_ (Millipore Sigma, 72060), 1X cOmplete™ Protease Inhibitor Cocktail (Millipore Sigma, 4693159001), and Sodium deoxycholate (Sigma-Aldrich, D6750-25G) on ice for 15 min. Lysates were centrifuged at 17,000 x g at 4 °C for 10 min, and the liquid phase was collected for use in downstream applications. Total protein content of the lysates was quantified (BCA Assay Kit, Thermo Fisher Scientific) and equalized among samples using the BCA method (Fisher Scientific, 23209). Samples were loaded in 7.5 or 10% TGX Stain-Free™ FastCast™ acrylamide gels (BioRad, 1610183). Transfer to PVDA membranes was realized using a TransBlot Turbo apparatus with the buffers and voltages recommended by the manufacturer. Membranes were blocked with 0.001% Poly(vinyl alcohol) (Sigma-Aldrich, 363065-25G) for 2 minutes. Membranes were incubated with primary and secondary antibody dilutions overnight and for 1 hour, respectively. Membranes were washed thrice in PBS 0.05% Tween-20 after incubation with primary and secondary dilutions. Images were captured with a Chemidoc system (BioRad) or using X-ray films.

Antibodies used (Supplier, Catalog#, Dilution factor):

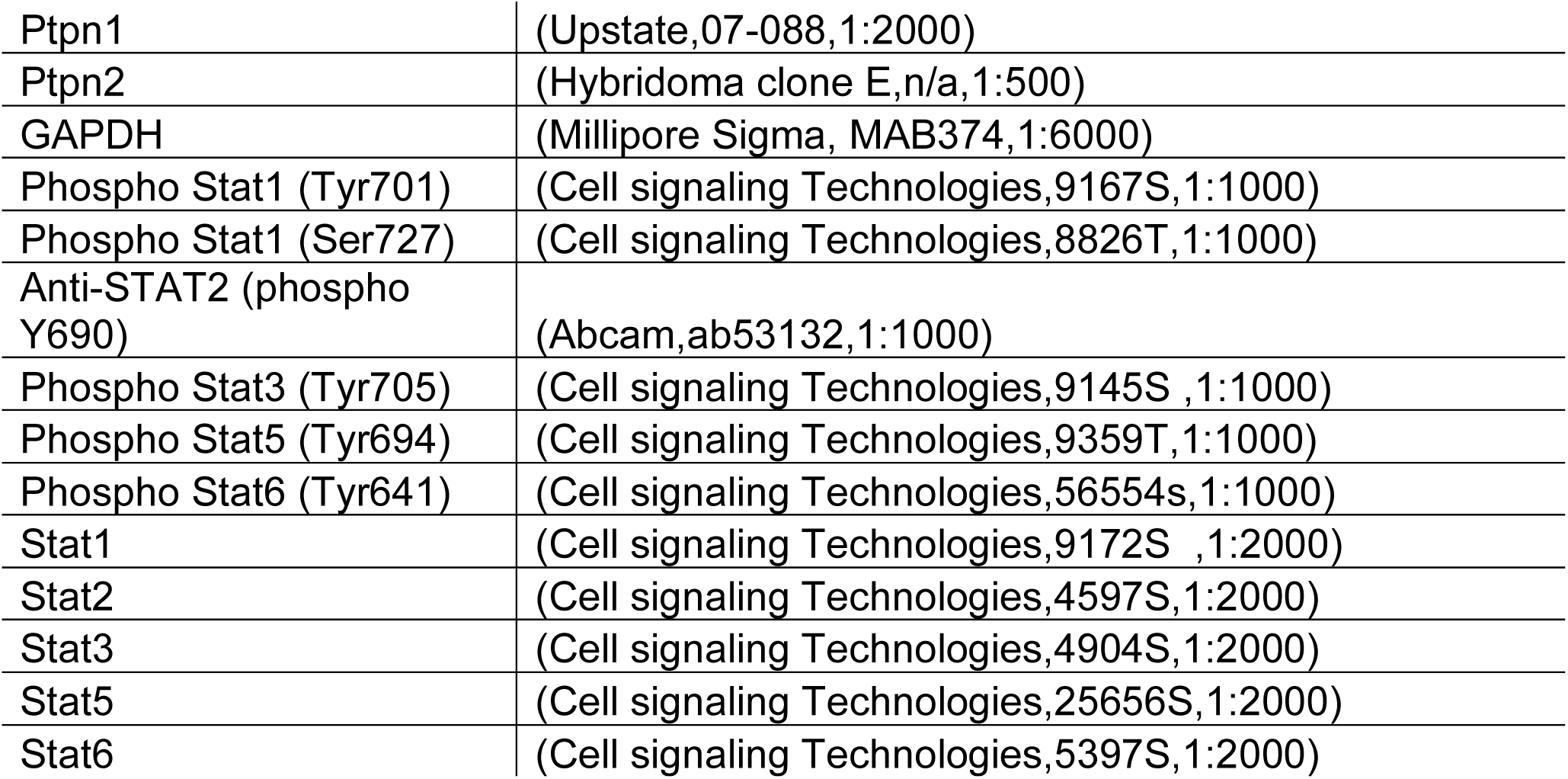

### Syngeneic tumor growth models in mice

Experiments on MC-38 (Colon cancer) and EMT6 (TNBC) tumors were realized by an independent CRO (Oncodesign, Montréal, Québec, Canada). For the colon cancer model, C57BL/6 mice were subcutaneously implanted with MC-38 tumor fragments obtained frozen from the Division of Cancer Treatment, Tumor Repository (National Cancer Institute, USA). For the TNBC breast cancer model, 1×10^6 EMT6 cells were injected subcutaneously in the right flank of BALB/c (BALB/cByJ) mice. Animals were randomized based on their individual tumor volume. Randomization was performed when the tumor on the right flank reached a mean of 75-150 mm^3^ (Days 14 and 11 respectively). Animals were randomized into four groups of ten animals each. Homogeneity between groups was tested by an analysis of variance (ANOVA). Control animals received an Intraperitoneal (IP) injection of control IgG (BioXcell, BE0089) at a dose of 10 mg/kg, twice a week for 3 consecutive weeks (TWx3). Group 2 animals received an Intraperitoneal (IP) injection of KQ-791 at a dose of 250 mg/kg, every two days for a total of ten injections (Q2Dx10). Group 3 animals received an Intraperitoneal (IP) injection of anti-PD1 (BioXcell, BE0146) at a dose of 10 mg/kg, twice a week for 3 consecutive weeks (TWx3). Group 4 animals received an Intraperitoneal (IP) injection of a combination of KQ791 at a dose of 250 mg/kg, every two days for a total of ten injections (Q2Dx10) and anti-PD1 at a dose of 10 mg/kg, twice a week for 3 consecutive weeks (TWx3). Mice were monitored every two days. For the tumor growth in LLC1-lung cancers, we injected 2 × 10^5^ LLC1 cells in 100μL of a solution containing equal parts Matrigel (Corning, 354262) and HBSS (Life Technologies, 14-025-092). Animals were injected with 500μL of a 250mg/kg dose of KQ791 and/or with 3.3mg/kg of anti-PD1 I.P. We realized a total of 7 injections of KQ791 and 5 injections of anti-PD1 over the course of the 21-day experiment. In subsequent experiments, we injected 2x10^5^ sgCTL or sgDKO B16-OVA, LLC1 or MC38-OVA cells in the right flank of C57BL/6 mice. Cells were also injected in 100μL of a solution containing equal parts Matrigel (Corning, 354262) and HBSS (Life Technologies, 14-025-092). For these experiments, we realized a total of 6 injections of KQ791 (Q2DX6) and/or 4 injections of anti-PD1 (TWX2), when applicable.

For the quantification of OVA_257-264_ Dextramer-positive peripheral Cd8+ T cells, we injected 2x10^5^ sgCTL or sgDKO B16-OVA cells in the right flank of C57BL/6 and treated the animals with 500μL of a 100mg/kg dose of KQ791, when applicable. KQ791 was injected every two days for a total of 4 injections starting day 5 post-tumor inoculation. Tissues (Spleen, tdLNs and Tumors) were collected at the experimental endpoint at day 15.

### Bone marrow Dendritic cell (BMDCs) differentiation and antigen cross-presentation assays

Bone marrow from both femurs and tibiae was harvested in RPMI supplemented with 10% heat-inactivated FBS (Life Technologies, 12483020). Marrows were harvested with centrifugation (10000xg for 30 seconds) and then filtered through a 70 μm Nylon cell strainer (Corning, 07-201-431) to remove solid fragments. BMDCs were generated during a 6-day differentiation protocol using 20/mL of recombinant murine GM-CSF (Preprotech, 315-03) and 20 ng/mL of recombinant murine IL-4 (Preprotech, 214-14). BMDCs were initially seeded at 5x10^5^ cells per mL in 12-well plates. KQ791 was added to the differentiating BMDCs on day 3 and kept at a concentration of 20μM until the end of the experiments. At the end of the differentiation period, BMDCs were harvested by pipetting up and down (Loosely adherent cells). For evaluation of antigen cross-presentation, BMDCs were inoculated with either 1μg/mL OVA_257-264_ peptide (Sigma-Aldrich, S7951) or 2x10^5^ B16-MO4 (sgCTL or sgDKO) for 24 hours. For co-culture experiments with OT-1 naïve CD8^+^ T cells, T cells, and BMDCs were co-cultured at a 1:1 ratio with each cell type at 1x10^5^ cells/well of a 12-well plate. OT-1 naïve T cells were isolated from the spleen using an immunomagnetic separation kit (StemCell Technologies, 18958). OT-1 naïve T cells were then labeled with CellTrace™ CFSE Cell Proliferation Kit (Life Technologies, C34554) as per the manufacturer’s instructions. OT-1 naïve T cells were then added to the OVA or B16 pulsed BMDCs and left to proliferate for 72h.

### Proliferation and LDH release assays

For proliferation curves (Incucyte), 8 replicate wells of B16, MC38, LLC1, or human cells were seeded at 2.5 × 10^3 to 5 × 10^3 cells/well in a 96-well plate and brought to the S3 Incucyte (Sartorius). 2 scans per well were scheduled for every 4 hours until confluency was reached. Average confluency at each timepoint was quantified using the Live-Cell Analysis System (Sartorius) with filter area >65μm2 and cleanup pixels -3. For measuring the impact of cytokine stimulation, B16 or LLC1 cells were seeded at 5x10^3^ cells per well of a 96-well plate and co-cultured with or without 20ng/mL of recombinant murine IFNβ1 (BioLegend, 581302), IFNγ (Preprotech, 315-05), and TNF-α (Preprotech, 315-01). Supernatants were collected and analysed for LDH release after 48 hours using the CytoTox 96® Non-Radioactive Cytotoxicity Assay kit (Promega, G1780). Cell density was quantified using the CyQUANT™ Cell Proliferation Assay Kit (Life Technologies, C7026).

## Statements

## Acknowledgements

We thank all members of the Tremblay laboratory and especially for their thorough overview of the manuscript. We are grateful for the support of R. Francoeur and A. Godille. We thank the Flow-cytometry and CMARC core facilities of the Life Science complex of McGill University. We highlight the work of R. Michaud, A. Laliberté, and C. Gagné for their care of the mice. We also highlight the work of Denis Faubert and Josée Champagne at the IRCM proteomics core facility and Lorne Taylor and Amy Wong from the RI-MUHC proteomics team.

## Author contributions

Conceptualization: A.P.

Methodology: A.P, E.W, R.S, I.A, C.W, E.S, S.U, A.H, B.C, S.H, S.D

Investigation: A.P, E.W, R.S, I.A

Visualization: A.P, E.W

Writing of the original draft: A.P.

Funding acquisition: M.L.T

Project administration: M.L.T

## Data and materials availability

The mass spectrometry proteomics data have been deposited in the ProteomeXchange Consortium (http://proteomecentral.proteomexchange.org) via the PRIDE partner repository with the dataset identifier PXD064419 (Project **DOI:** 10.6019/PXD064419). The transcriptomic dataset has been deposited as a GEO dataset: GEO accession GSE297993. For reviewer access, use the following link and token: Go to https://www.ncbi.nlm.nih.gov/geo/query/acc.cgi?acc=GSE297993. Enter the token khslmecebpavfwz into the box. All other data needed to evaluate the conclusions in the paper are present in the paper or the Supplementary Materials.

---

**Extended Data Fig. 1:**
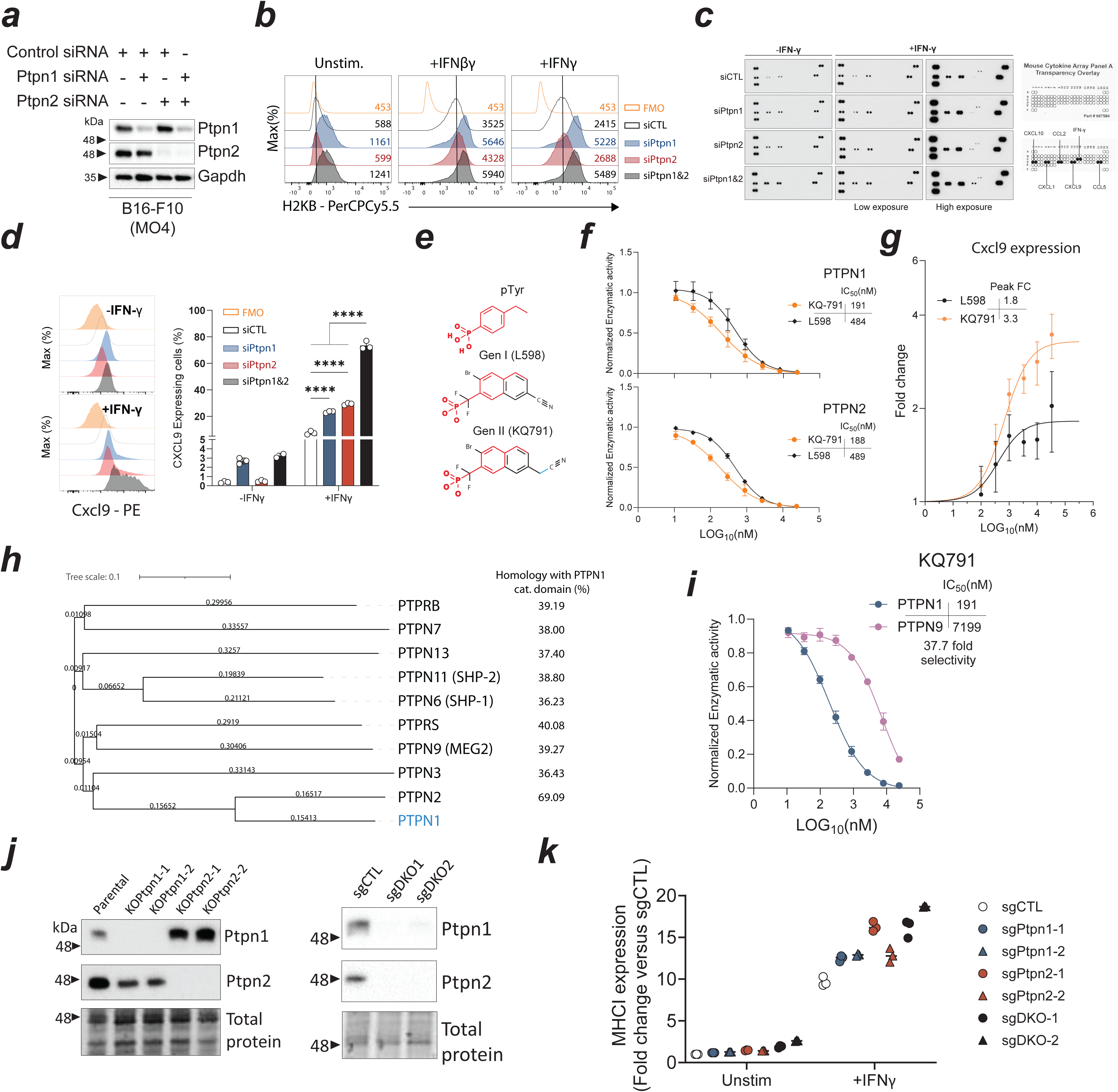
Supporting data for Fig. 1. a) Western blotting for Pptn1 and Ptpn2 in B16-OVA cells transfected with the indicated siRNAs. Gapdh is used as the loading control. b) Flow cytometric analysis of H2KB (MCHI) expression in B16-OVA cells after 24h of stimulation with IFNβ or IFNγ (20ng/mL). FMO: Fluorescence Minus One staining control. c) Raw dot blot of the mouse cytokine/chemokine array membranes with or without stimulation with IFNγ for 24h. Low (30 sec) and high exposure (5 min) pictures were included to detect cytokines/chemokines with lower abundance. d) Histograms (left) and bar chart (right) of Cxcl9 expression in B16-OVA cells as measured by flow cytometry. e) Molecular structures of phospho-tyrosine (pTyr), PTPN1/2 dual inhibitor lead (Gen I, L598), and KQ791 (Gen II). f) In vitro phosphatase activity curves (IC_50_) for PTPN1 and PTPN2 with increasing concentrations of L598 and KQ791. g) Flow cytometric analysis of Cxcl9 production in B16-OVA cells treated with IFNγ (24h) and with increasing concentrations of L598 or KQ791. Data were normalized to the mean fluorescence intensity (MFI) of cells treated with IFNγ alone h) Phylogenic tree of the phosphatases with significant structural similarity to PTPN1. The catalytic domain of PTPN1 (Amino acids 3-277) was used for protein-BLAST alignment. i) In vitro phosphatase activity curves (IC_50_) for PTPN1 and PTPN9 with increasing concentrations of KQ791. j) Western blotting for Pptn1 and Ptpn2 in B16-OVA cells that were subjected to CRISPR/Cas9 mediated knockout of Ptpn1 or Ptpn2. Total protein is used as the loading control (left). Western blotting for Pptn1 and Ptpn2 in B16-OVA cells subjected to double CRISPR/Cas9-mediated knockout (right). k) Flow cytometric analysis of MHCI (H2KB) expression in two distinct CRISPR/Cas9 knockout pools for each indicated condition. Data was normalized to unstimulated control guide RNA (sgCTL) cells. Data representative of at least three independent experiments (a, b, d, f, g). Error bars represent mean±s.e.m. Statistical analysis: Two-way ANOVA with Tukey’s multiple comparison test (d). Key: ns nonsignificant, *p<0.05, **p<0.01, ***p<0.001 and ****p<0.0001.

**Extended Data Fig. 2:**
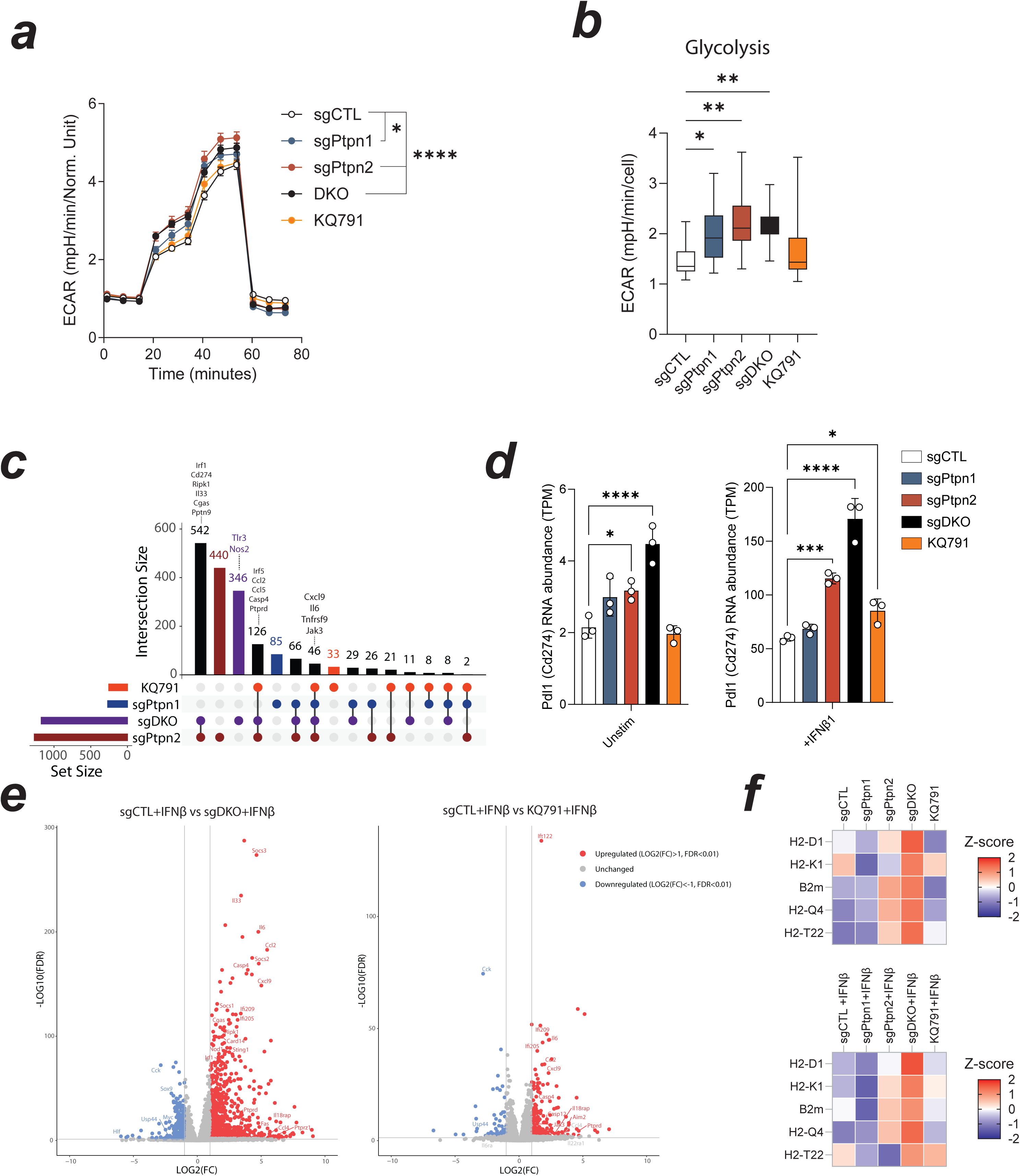
Metabolic and transcriptomic analysis of Ptpn1/2 deficient B16-OVA cells. a) Extracellular Acidification Rate (ECAR) assay in B16-OVA cells with the indicated knockout or treatments as measured by mpH/min/cell change. Data was normalized to cell content per well. Glucose was added 20 minutes after plate incubation. Oligomycin was added 40 minutes after plate incubation. b) Mean ECAR during the glycolysis phase of ECAR measurement (20-40 minutes). c) Upset plot of the differentially expressed genes (DEGs) that are common between the indicated sets of conditions (6h post-IFNβ treatment). Genes of interest are labeled above the respective conditions that share these DEGs. d) Mean transcript per million (TPM) RNA abundance for Pdl1 (Cd274) in unstimulated and IFNβ-stimulated B16-OVA cells with the indicated conditions. e) Volcano plots for the DEGs in IFNβ-stimulated sgDKO (left) and IFNβ-KQ791-treated (right) compared to IFNβ-stimulated sgCTL B16-OVA cells (LOG2(FC)>1 or <-1, FDR<0.01). f) Z-score heatmap for MHCI-related genes in unstimulated (top) or IFNβ-stimulated (bottom) B16-OVA cells. Error bars represent mean±s.e.m. Statistical analysis: One-way ANOVA with Dunnett’s multiple comparison test (b and d). Key: ns nonsignificant, *p<0.05, **p<0.01, ***p<0.001 and ****p<0.0001.

**Extended Data Fig. 3:**
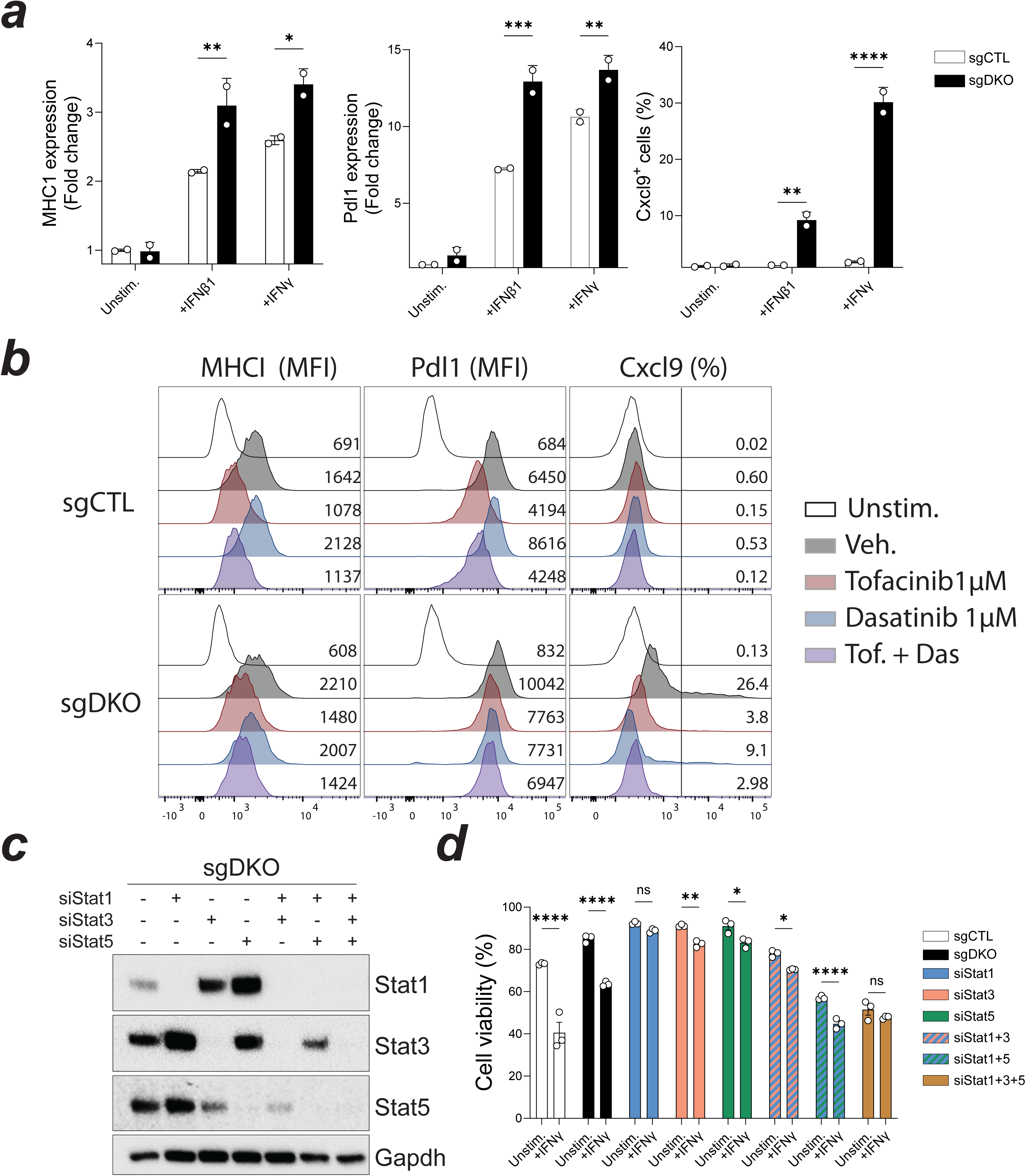
Supporting data for Fig. 3. a) Flow cytometric analysis of MHCI (H2KB), Pdl1 and Cxcl9 expression fold-change in B16-OVA cells stimulated with IFNβ or IFNγ. Data was normalized to unstimulated sgCTL cells. b) MHCI (H2KB), Pdl1, and Cxcl9 expression histograms in B16-OVA sgDKO cells treated with IFNγ and the indicated drugs for 24 hours. Unstimulated sgDKO (no IFNγ) are used as the negative staining control. c) Western blotting for Stat1, Stat3 and Stat5 in B16-OVA sgDKO cells transfected with the indicated siRNAs. Gapdh is used as the loading control. d) Bar chart of the frequency of viable B16-OVA sgCTL or sgDKO cells transfected with the indicated siRNAs after 24hours of stimulation with IFNγ. Viability was assessed by flow cytometry using a fixable viability dye (eFluor780). Data representative of at least three independent experiments. Error bars represent mean±s.e.m. Statistical analysis: Two-way ANOVA with Šídák’s multiple comparisons test (a and d). Key: ns nonsignificant, *p<0.05, **p<0.01, ***p<0.001 and ****p<0.0001.

**Extended Data Fig. 4:**
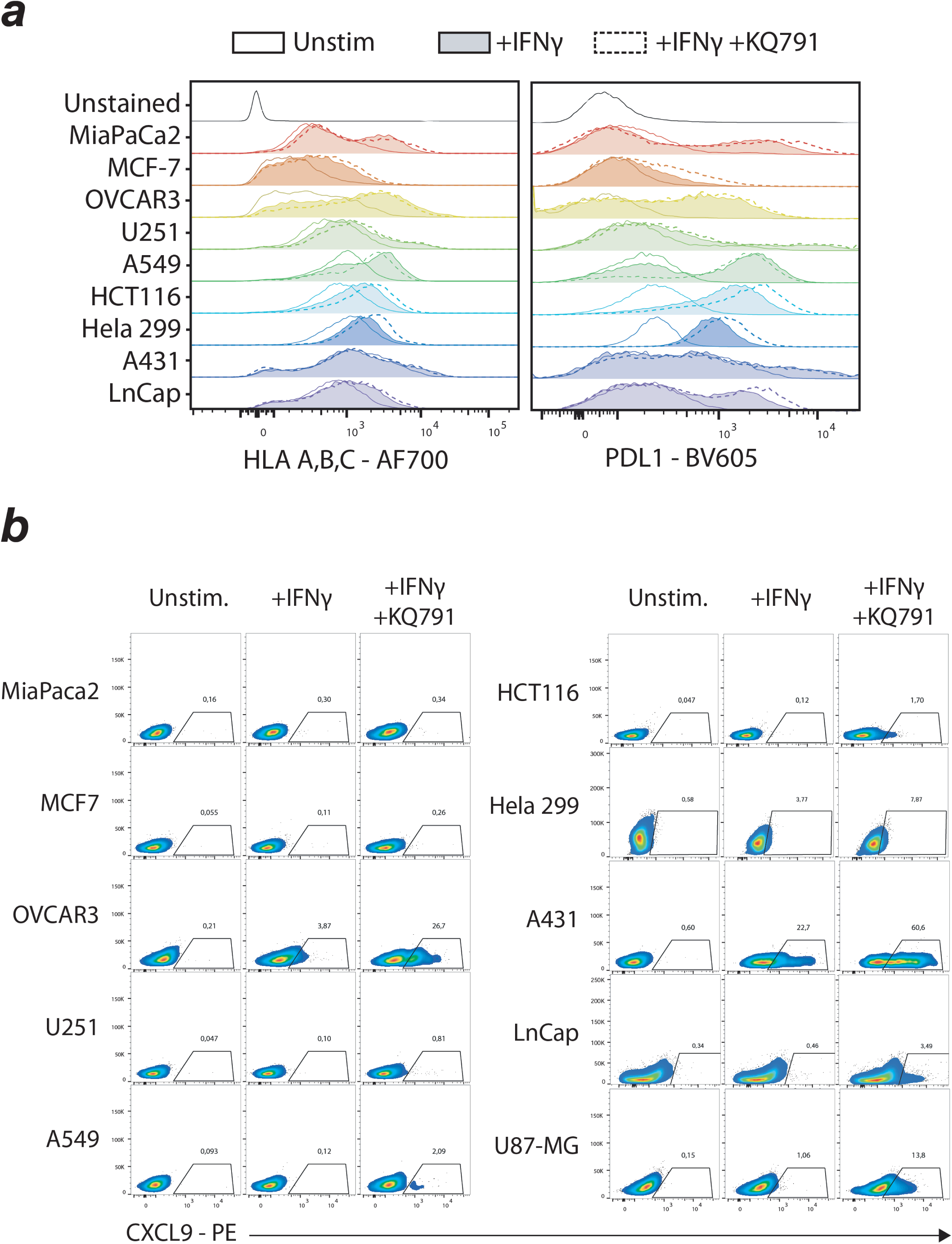
Supporting data for Fig. 4. a) HLA A, B, C (MHCI) and PDL1 expression histograms in human cancer cell lines as measured by flow cytometry 24h after IFNγ stimulation. b) Flow cytometric analysis of CXCL9 expression in human cancer cell lines. Data representative of three independent experiments done in triplicate.

**Extended Data Fig. 5:**
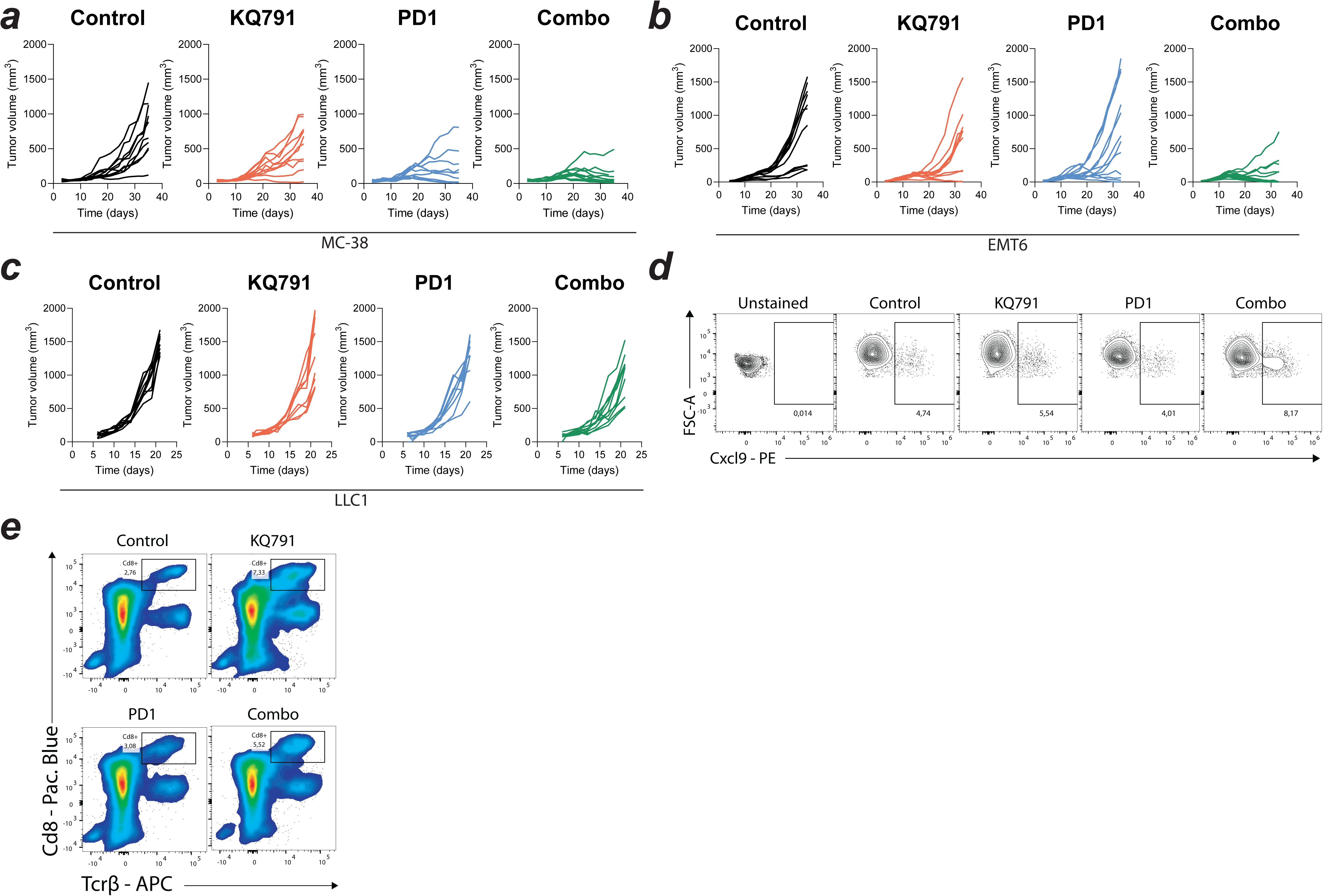
Supporting data for Fig. 5. a) Individual tumor growth curves in MC38 cancer bearing mice treated with KQ791(red), anti-PD1 (blue), a combination of both (combo, green), or with an isotype control (black). Treatment regimens: KQ791 Q2DX10. anti-PD1 or control TWx3. N=10/group. b) Individual tumor growth curves in EMT6 cancer bearing mice treated with KQ791(red), anti-PD1 (blue), a combination of both (combo, green), or with an isotype control (black). Treatment regimens: KQ791 Q2DX10. anti-PD1 or control TWx3. N=12/group. c) Individual tumor growth curves of LLC1 lung cancer bearing mice treated with KQ791, anti-PD1, a combination of both (combo), or with an isotype control. Treatment regimens: KQ791 Q2DX7, anti-PD1 TWx3. N=10/group. d) Contour plot of Cxcl9 expression in LLC1 cancer cells as measured by flow cytometry. e) Flow cytometric analysis of Cd8^+^ T cell frequency within the Cd45^+^ compartment (Cd8^+^, Tcrβ^+^). Gated on live Cd45^+^ cells from LLC1 tumors.

**Extended Data Fig. 6:**
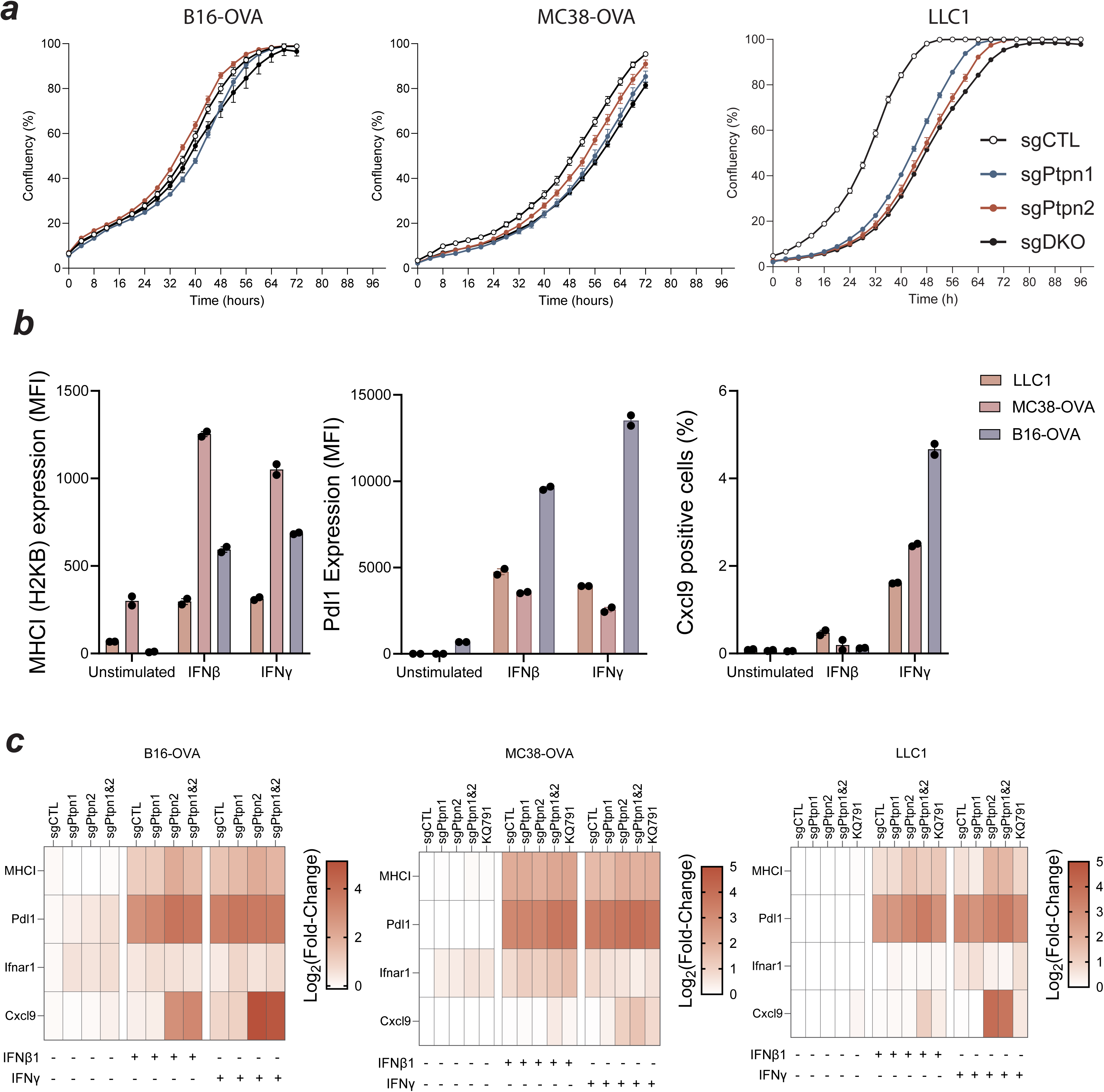
Validation of the intrinsic role of Ptpn1/2 in MC38-OVA and LLC1 cancer cell lines. a) Cell proliferation curves for B16-OVA, MC38-OVA and LLC1 cancer cells at steady state. b) Bar chart of MHCI (H2KB), Pdl1, and Cxcl9 expression fold-change in B16-OVA, MC38-OVA, and LLC1 cancer cells after stimulation with IFNβ or IFNγ (24h). c) Log2(Fold-change) heatmap of MHCI (H2KB), Pdl1, Ifnar1 and Cxcl9 expression in B16-OVA, MC38-OVA and LLC1 cells with the various Ptpn1/2 genotypes. Data was normalized to unstimulated sgCTL cells within each cell line and marker. Data representative of three independent experiments done in duplicates or triplicate. Error bars represent mean±s.e.m.

**Extended Data Fig. 7:**
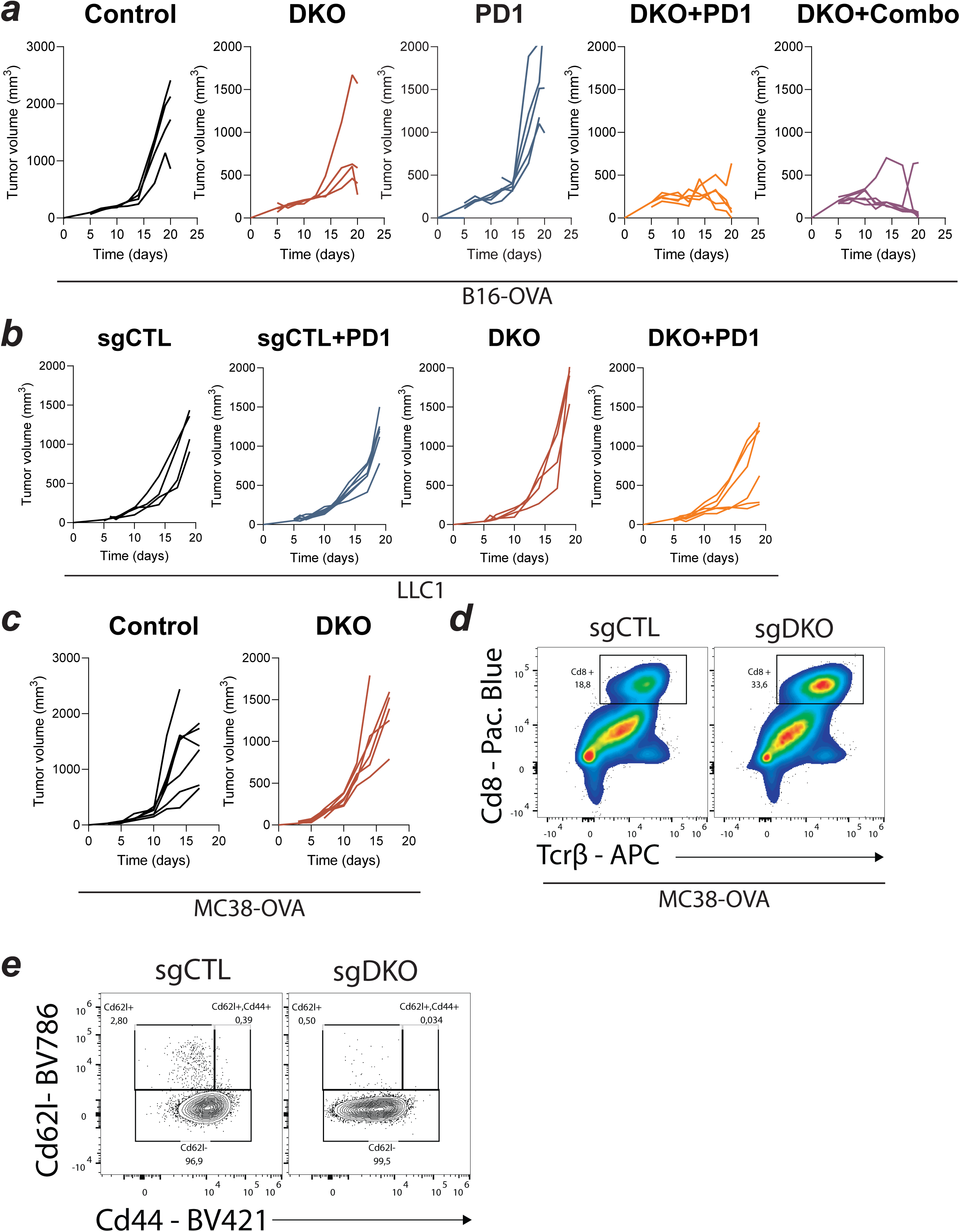
Supporting data for Fig. 6. a) Individual tumor growth curves of sgCTL or sgDKO B16-OVA melanomas treated with anti-PD1, anti-PD1 and KQ791 (Combo), or left untreated (control). Treatment regimens: anti-PD1 TWx2. KQ791 Q2DX6. N=4-6/group. b) Individual tumor growth curves of sgCTL or sgDKO LLC1 lung cancers treated with anti-PD1, anti-PD1 or left untreated (control). Treatment regimens: anti-PD1 TWx2. KQ791 Q2DX6. N=4-6/group. c) Individual tumor growth curves of sgCTL or sgDKO MC38-OVA cancers (no treatment). N=6/group. d) Flow cytometric analysis of Cd8^+^ T cell frequency within the Cd45^+^ compartment (Cd8^+^, Tcrβ^+^). Gated on live Cd45^+^ cells from MC38-OVA sgCTL or sgDKO tumors. e) Contour plot of Cd62l^+^, Cd62l^-^, and Cd44^+^ populations within Cd8^+^ T cells of sgCTL or sgDKO MC38-OVA tumors as measured by flow cytometry. Gated on live Cd45^+^, Tcrβ^+^, Cd8^+^.

**Extended Data Fig. 8:**
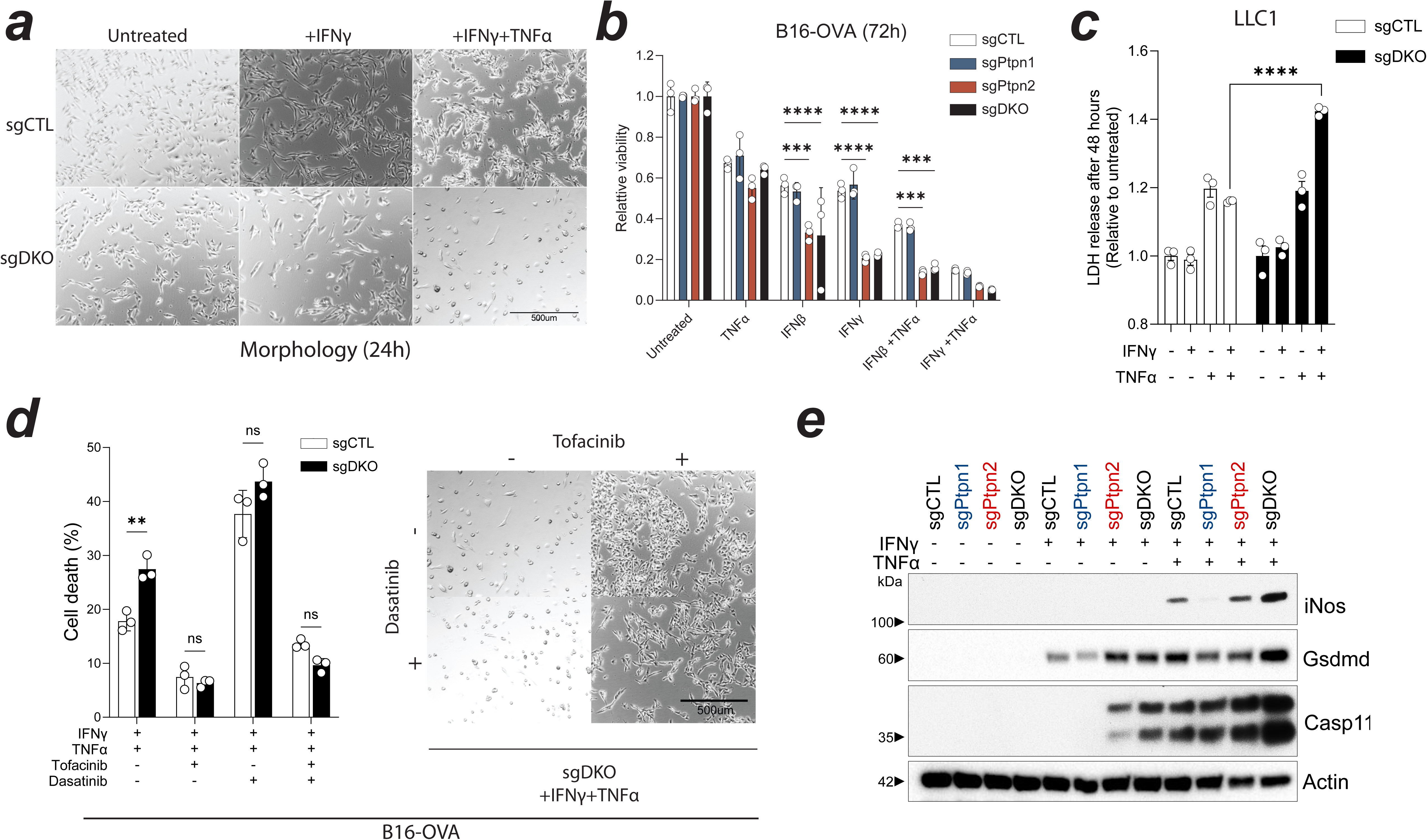
Supporting data for Fig. 7. a) Micrograph of sgCTL or sgDKO B16-OVA cells 24 hours post-treatment with IFNγ or IFNγ and TNFα. The scale bar represents 500μm. b) Relative cell abundance/viability of sgCTL, sgPtpn1, sgPtpn2, and sgDKO B16-OVA cells cultured with IFNβ, IFNγ, and/or TNFα for 72h. Data was normalized to the untreated cells in each genotype. c) Relative LDH release in sgCTL and sgDKO LLC1 cultures with IFNγ, and/or TNFα (48h post-stimulation). Data was normalized to the untreated cells in each genotype. d) Frequency of dead cells in sgCTL or sgDKO B16-OVA cells 36 hours post-stimulation with IFNγ & TNFα and post-treatment with Tofacitinib and/or Dasatinib (1μM). Dead cells were measured with flow cytometry using a fixable live/dead stain (eFluor780^+^ cells). Micrograph of sgDKO cells 24h post-stimulation with IFNγ & TNFα and treatment with or without Tofacitinib and/or Dasatinib (1μM) (right). The scale bar represents 500μm. e) Western blotting of iNOs, Gasdermin D (Gsdmd) and Caspase 11 in sgCTL, sgPtpn1, sgPtpn2, and sgDKO B16-OVA cells cultured with IFNγ or IFNγ + TNFα for 36 hours. Actin is used as the loading control. Data representative of at least three independent experiments done in triplicate. Error bars represent mean±s.e.m. Statistical analysis: Two-way ANOVA with Dunnett’s multiple comparison test (b and c). Two-way ANOVA with Tukey’s multiple comparison test (d). Key: ns nonsignificant, *p<0.05, **p<0.01, ***p<0.001, and ****p<0.0001.

**Extended Data Fig. 9:**
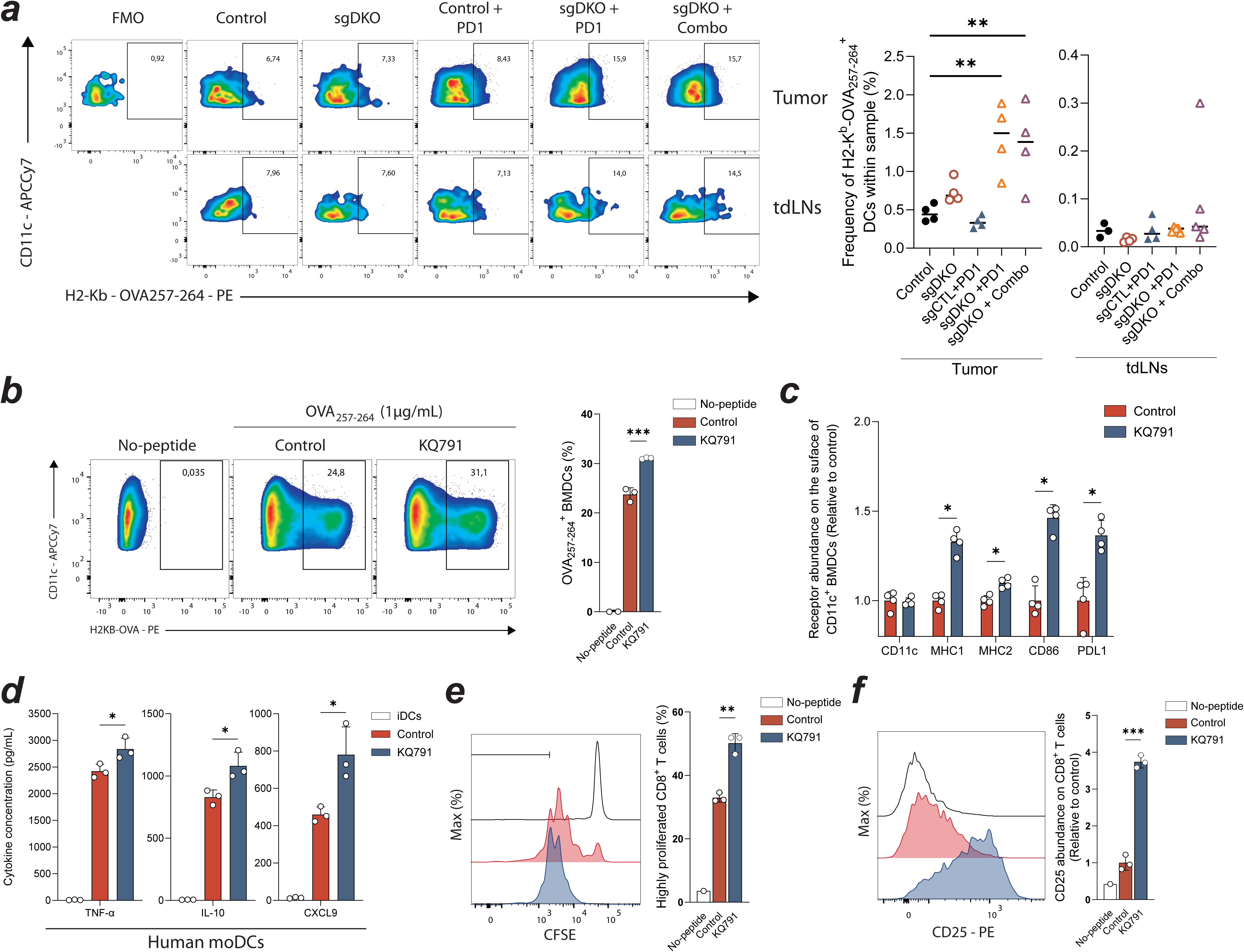
Ptpn1/2 inhibition enhances the functionality and antigen cross-presentation capabilities of dendritic cells. a) Flow cytometric analysis of the frequency of dendritic cells with MHCI (H2KB) bound to OVA_257-264_ in tumor (top) or tumor draining lymph nodes (tdLNs, bottom) of mice bearing B16-OVA tumors (sgCTL or sgDKO treated with PD1 or combo). N=4/group. b) Frequency of MHCI (H2KB) bound to OVA_257-264_ in bone marrow-derived dendritic cells (BMDCs) as measured by flow cytometry. c) Flow cytometric analysis of Cd11c, MHCI (H2KB), MHC2, Cd86 and Pdl1 in BMDCs treated with or without KQ791 during their differentiation. Mean fluorescence intensity (MFI) was normalized to the control BMDCs for each marker. d) Cytokine multiplexing in mature human monocyte-derived dendritic cells (moDCs) treated with or without KQ791 during their differentiation. TNF-α, IL-10, and CXCL9 were the factors with the most significant increase in abundance after KQ791 treatment among a panel of 48 cytokines. e) CFSE proliferation histograms of OT-1 T cells co-cultured with KQ791-treated or untreated BMDCs pulsed with OVA_257-264_ for 72h (left). Bar graph of the frequency of highly proliferated Cd8^+^ T cells from the CFSE analysis (CFSE MFI<2x10^3^) (right). f) Cd25 expression histograms in OT-1 T cells co-cultured with KQ791-treated or untreated BMDCs pulsed with OVA_257-264_ for 72h (left). Bar graph of the relative abundance of Cd25 in OT-1 Cd8^+^ T cells (right). Data was normalized to the control, untreated OVA_257-264_ pulsed BMDCs, stimulated T cells (red). Data representative of at least three independent experiments done in triplicate or quadruplicate (b-f). Error bars represent mean±s.e.m. Statistical analysis: One-way ANOVA with Dunnett’s multiple comparison test (a, e, and f). Multiple unpaired T-tests (c and d). Key: ns nonsignificant, *p<0.05, **p<0.01, ***p<0.001, and ****p<0.0001.

**Extended Data Fig. 10:**
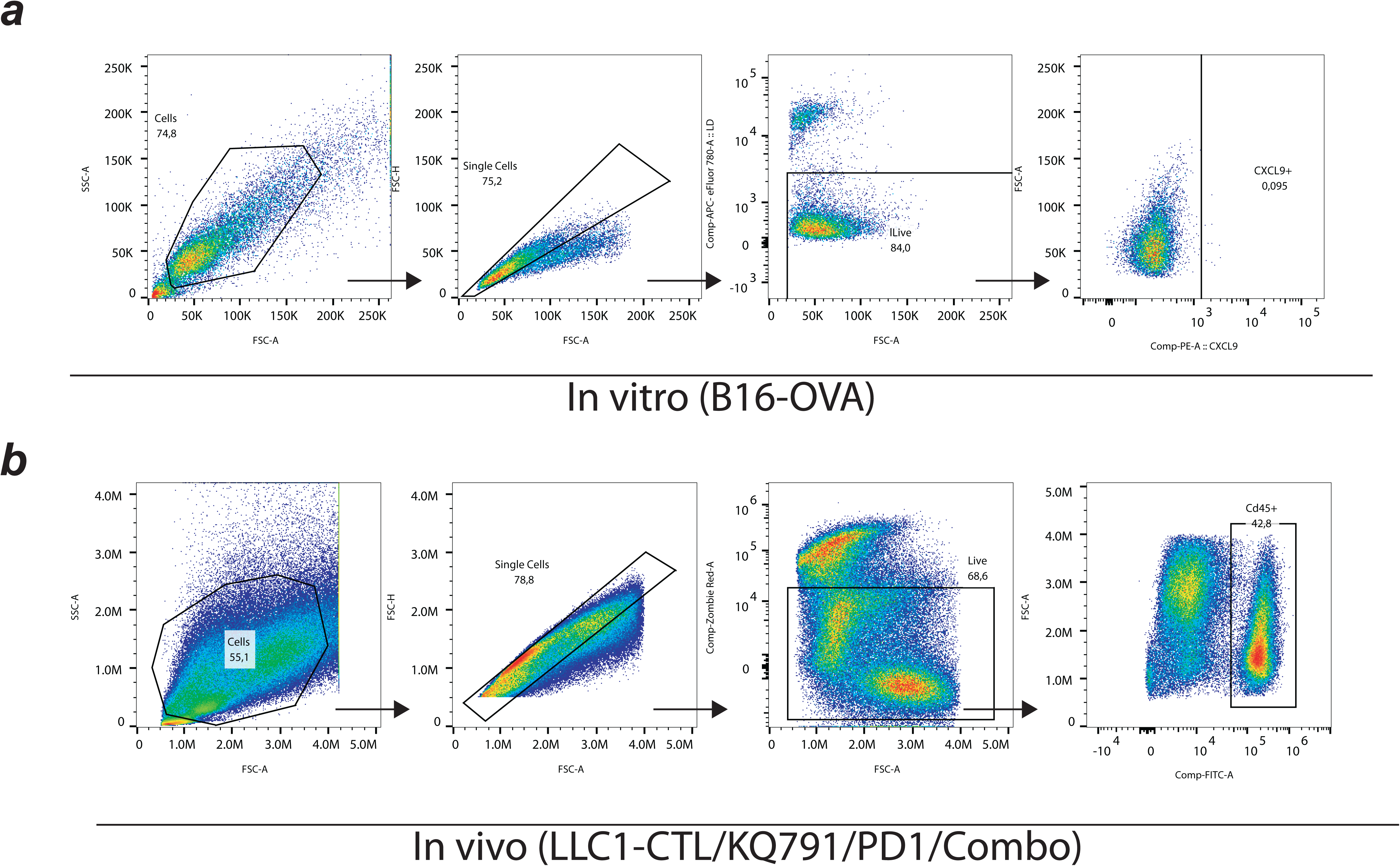
Gating strategies for in vitro and in vivo studies. a) From left to right: Gating strategies for quantifying MHCI (H2KB), Pdl1, and Cxcl9 expression in in vitro cultures. b) From left to right: Gating strategies for assessing immune cell composition in tumors from in vivo experiments.

